# Timing of immune checkpoint blockade shapes anti-tumor immunity via a clock-dependent chemokine axis

**DOI:** 10.1101/2025.11.03.686293

**Authors:** Jake N. Lichterman, Tarun Srinivasan, Ruheng Wang, Chaitanya Dende, Christina Zarek, Josue I. Ramirez, Noheon Park, Hesuiyuan Wang, Wenling Li, Jiwoong Kim, Laura Coughlin, Nicole Poulides, Parastoo Sabaeifard, Suzette N. Palmer, Aidan C. Morrill, Brian Hassell, John M. Shelton, Xin V. Li, Bret M. Evers, Xiaowei Zhan, Joseph S. Takahashi, Lora V. Hooper, Andrew Y. Koh

## Abstract

Circadian clocks regulate immunity, yet how they shape the tumor immune microenvironment and influence cancer immunotherapy remains unclear. Here, we show that tumor immune infiltration and immune checkpoint inhibitor efficacy vary by time of day in mice, driven by intrinsic clocks in dendritic cells and CD8^+^ T cells. Time-of-day modulates the abundance, spatial organization, and cytokine–chemokine production of tumor-infiltrating immune cells. Mechanistically, dendritic cell clocks control expression of *Cx3cl1*, driving recruitment of CX3CR1^+^CD8^+^ T cells and thereby reshaping the tumor immune microenvironment to enhance immunotherapy efficacy. Disruption of this axis abolishes time-of-day-dependent differences in treatment response. These findings identify a circadian mechanism of immune cell recruitment to tumors and provide mechanistic insight into clinical observations linking treatment timing to immunotherapy outcomes.

**One sentence summary:** Time of day determines cancer immunotherapy efficacy through a circadian clock–dependent chemokine axis.

## Main Text

Immune checkpoint inhibitor therapy (ICT) targeting PD-1/PD-L1 or CTLA-4 has revolutionized cancer treatment, yielding durable remission, and even cures, for select patients with previously incurable cancers (*1–5*). Most patients, however, do not achieve lasting clinical responses. The mechanisms that regulate sustained anti-tumor immunity remain poorly understood.

The tumor immune microenvironment (TIME) is a critical determinant of ICT efficacy. Tumors enriched with effector CD8^+^ T cells and activated myeloid cells, particularly conventional dendritic cells (cDCs), respond more favorably to ICT than tumors with sparse or dysfunctional immune cell infiltrates (*6–8*).

Host-derived, tumor cell–extrinsic factors have recently emerged as key modulators of the TIME and ICT response (*9, 10*). One such host factor, the circadian clock, remains largely unexplored in the context of cancer immunotherapy. The circadian clock is a conserved transcriptional–translational feedback loop that orchestrates daily rhythms in cellular function across nearly all mammalian cells (*11*). Circadian regulation of innate and adaptive immunity produces marked time-of-day fluctuations in leukocyte development, trafficking, antigen presentation, and cytokine production across the 24-hour cycle (*12, 13*). These fluctuations result in time-of-day-dependent responses to vaccines, pathogens, and tumor engraftment (*14–17*). Recent retrospective clinical studies report survival differences based solely on the time of ICT administration (*18-20*). While these observations suggest that ICT treatment timing matters, the underlying biological mechanisms remain unclear.

Here, we show that circadian regulation of key immune cells drives time-of-day differences in the composition and function of the TIME, ultimately shaping ICT efficacy. In preclinical cancer models, ICT is most effective when administered at the time of peak immune activity within the TIME, characterized by increased infiltration and activation of dendritic cells (DCs) and CD8^+^ T cells. Deletion of the core clock gene *Bmal1* in either immune cell type blunts treatment efficacy, demonstrating that ICT response depends on immune cell-intrinsic circadian clocks. We further identify a clock-controlled chemokine axis—CX3CL1–CX3CR1 signaling—that governs time-of-day-dependent immune cell recruitment. Circadian regulation of *Cx3cl1* in DCs drives CX3CR1^+^ CD8^+^ T cell recruitment into tumors, reshaping the TIME and enhancing ICT efficacy. These findings define a circadian mechanism that regulates tumor immune cell infiltration and suggest treatment timing as a critical, yet underappreciated, determinant of immunotherapy response.

### ICT efficacy varies by time of day of treatment across tumor models

Clinical studies suggest that immunotherapy outcomes may depend on the time of ICT administration (*18–20*). To test this, we evaluated whether ICT efficacy is time-dependent in mice. We used the well-established MC38 adenocarcinoma model (*21, 22*) and administered ICT (anti– PD-1 monoclonal antibody or isotype control) at six zeitgeber times (ZT): ZT2, ZT6, ZT10, ZT14, ZT18, and ZT22. ZT refers to hours after light onset in a 12-hour light-dark cycle (Fig. 1A). Mice are inactive during ZT0–ZT12 (light phase) and active during ZT12–ZT24 (dark phase). ICT administered at ZT2 (two hours into the light/inactive phase) resulted in smaller tumor volumes (Fig. 1B and C), lower tumor weights (Fig. S1A), and increased survival time (Fig. 1D) compared to treatment at ZT18 (six hours into the dark/active phase). In contrast, isotype-treated mice showed no time-of-day differences in tumor growth, suggesting that this effect is specific to ICT (Fig. S1B).

**Figure 1.**
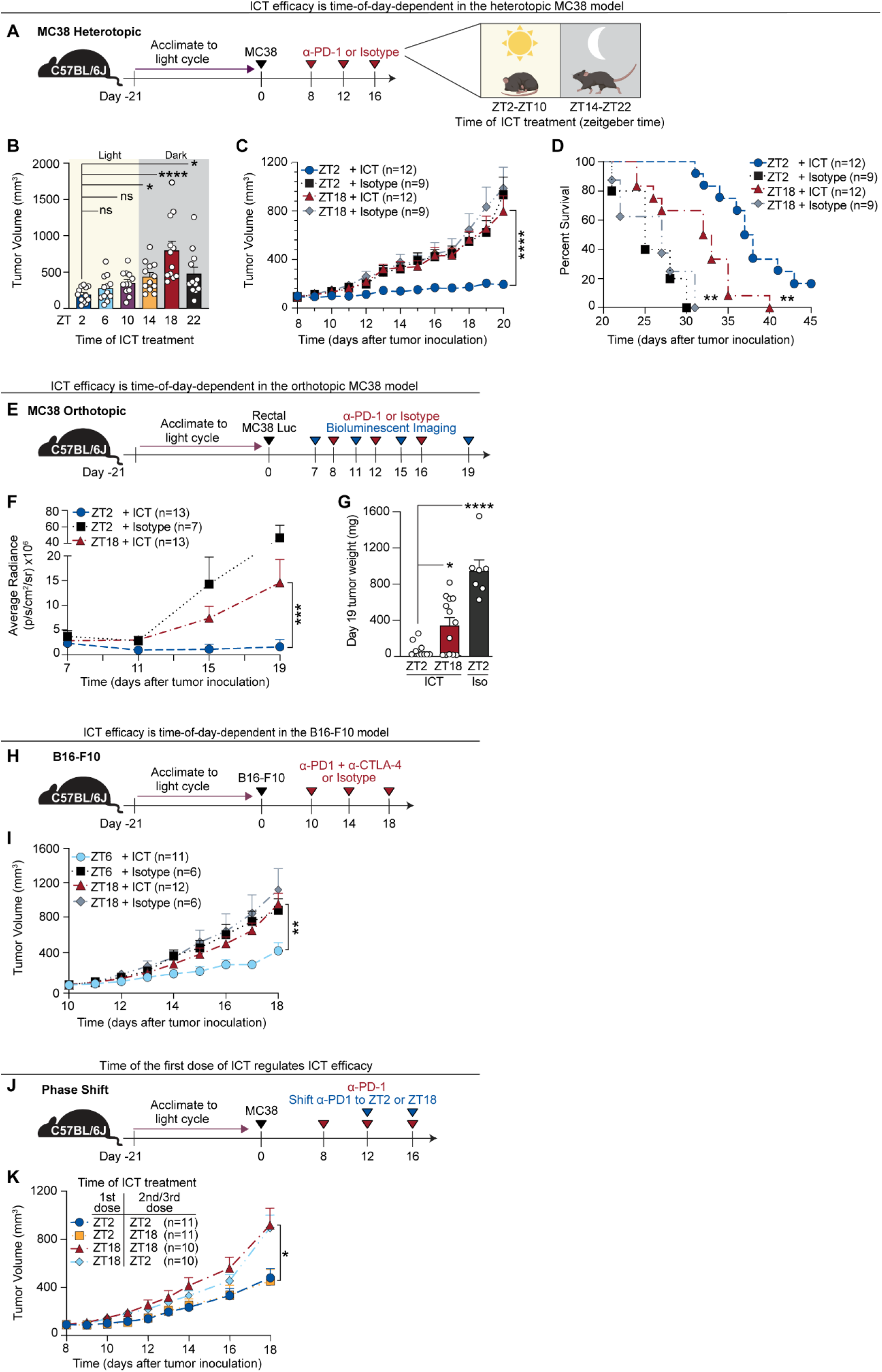
ICT efficacy varies by time of day of treatment across tumor models. Three syngeneic tumor models were used to assess the impact of ICT treatment time on ICT efficacy. All experiments were performed in 8-week-old female C57BL/6J mice acclimated to a 12-hour light/dark cycle for 21 days prior to tumor inoculation at ZT3. **(A)** Schematic overview of the MC38 heterotopic model. MC38 cells were injected subcutaneously into the right inguinal flank. Mice received 200 µg of anti–PD-1 (ICT) or isotype control (rat IgG2a) intraperitoneally every 4 days for three doses, beginning when tumor volumes reached 100 ± 30 mm^3^. **(B)** Tumor volumes at day 20 following ICT administered at six distinct zeitgeber times (ZT2– ZT22); n = 12 mice per group. **(C)** Tumor volumes following ICT or isotype treatment at ZT2 or ZT18; n = 9–12 mice per group. **(D)** Survival of tumor-bearing mice described in (C); n = 9–12 mice per group. **(E)** Schematic of orthotopic MC38-luciferase (MC38-Luc) colorectal tumor model. MC38-Luc cells were implanted into the rectal lumen. Rectal MC38 tumors were imaged prior to ICT treatment on day 7 and mice were stratified into different groups with equivalent average bioluminescence. Mice were then treated with ICT (anti–PD-1) or isotype intraperitoneally every 4 days for three doses. Tumor response was monitored by bioluminescent imaging prior to each dose of ICT and final tumor weights were quantified on day 19. **(F)** Average bioluminescent radiance over time (p/s/cm^2^/sr) for MC38 tumor-bearing mice treated with ICT or isotype at ZT2 vs. ZT18. n = 7–13 mice per group. **(G)** MC38-Luc rectal tumor weights on day 19 after ICT or isotype at ZT2 vs. ZT18; n = 7–13 mice per group. **(H)** Schematic overview of the B16-F10 melanoma model. B16-F10 cells were injected subcutaneously into the right flank. Mice were treated with 200 µg anti–PD-1 and 200 µg anti– CTLA-4 antibodies or isotype controls every 4 days for three doses, starting on day 10, when tumor volumes reached 100 ± 30 mm^3^. **(I)** Tumor volumes from mice described in (H); n = 6–12 mice per group. **(J)** Schematic overview of the phase-shift experiment. Utilizing the heterotopic MC38 model as in (A), tumor-bearing mice received their first dose of anti–PD-1 at ZT2 or ZT18 on day 8, followed by two additional doses of anti–PD-1 on day 12 and 16 at either the same ZT (e.g. ZT2→ZT2) or the alternative ZT (e.g. ZT2→ZT18). **(K)** Tumor volumes from mice described in (J); n = 10–11 mice per group. ZT, zeitgeber time; ICT, immune checkpoint inhibitor therapy. Statistical analysis by Kruskal– Wallis test for (B, G), Mann-Whitney test for (C, F, I, K) or Log-rank test for (D). Points represent individual mice. Bars denote mean ± SEM. All results are representative of ≥2 independent experiments. *p < 0.05; **p < 0.005; ***p < 0.001; ****p < 0.0001; ns, not significant.

To determine whether the time-of-day effects on ICT efficacy extend to different tumor models, we tested two additional systems. First, we used an orthotopic colorectal tumor model by implanting luciferase-expressing MC38 cells into the rectum (Fig. 1E). As in the MC38 subcutaneous setting, ZT2 treatment reduced tumor burden, as evidenced by lower bioluminescence (Fig. 1F; Fig. S1C) and tumor weight (Fig. 1G) compared to ZT18. We next examined the B16-F10 melanoma model which is immunologically distinct, characterized by sparse immune cell infiltration in the tumor and relative resistance to anti-PD-1 monotherapy (*21-24*). Hence, mice were treated with combined anti-PD-1/CTLA-4 at ZT6 or ZT18 (Fig. 1H). Again, treatment during the inactive phase (ZT6) improved tumor control (Fig. 1I), suggesting that time-of-day-dependent ICT efficacy is preserved across tumor types and anatomical sites.

ICT antibodies (including anti–PD-1, anti–PD-L1, and anti–CTLA-4) exhibit long half-lives. In mice, terminal half-lives often range from several days to over a week; in humans, they can extend to weeks (e.g., ∼27 days for anti-PD-1 nivolumab) (*25–27*). Despite this prolonged pharmacokinetics, we observed marked differences in ICT efficacy with only a few hours’ difference in dosing time. Thus, we hypothesized that the time of the initial ICT dose may be sufficient to program a durable anti-tumor response. To test this, we performed a “phase-shift” experiment: mice received their first ICT dose at either ZT2 or ZT18, followed by two additional doses at either the same (e.g. ZT2→ZT2) or opposite time points (e.g. ZT2→ZT18) (Fig. 1J). Tumor control was determined by the timing of the first dose, regardless of when subsequent doses were given (Fig. 1K). These findings suggest that time-of-day differences present at the time of ICT initiation may imprint long-lasting effects on anti-tumor immunity.

### Time-of-day influences the spatial and cellular organization of the tumor immune microenvironment

To determine whether time-of-day differences in ICT efficacy arise from time-of-day variation in the TIME, we performed whole-transcriptome spatial profiling (Visium HD) on subcutaneous MC38 tumors harvested at ZT2 and ZT18 in ICT-naïve mice (Fig. 2A). Unsupervised clustering of spatial transcriptomic data identified both immune and non-immune cell populations, annotated based on canonical marker gene expression (Fig. S2A) and histomorphologic features (Fig. S2B). Indeed, tumors collected at ZT2 showed greater infiltration of myeloid and T cells compared to ZT18 (Fig. 2B; Fig. S2C).

**Figure 2.**
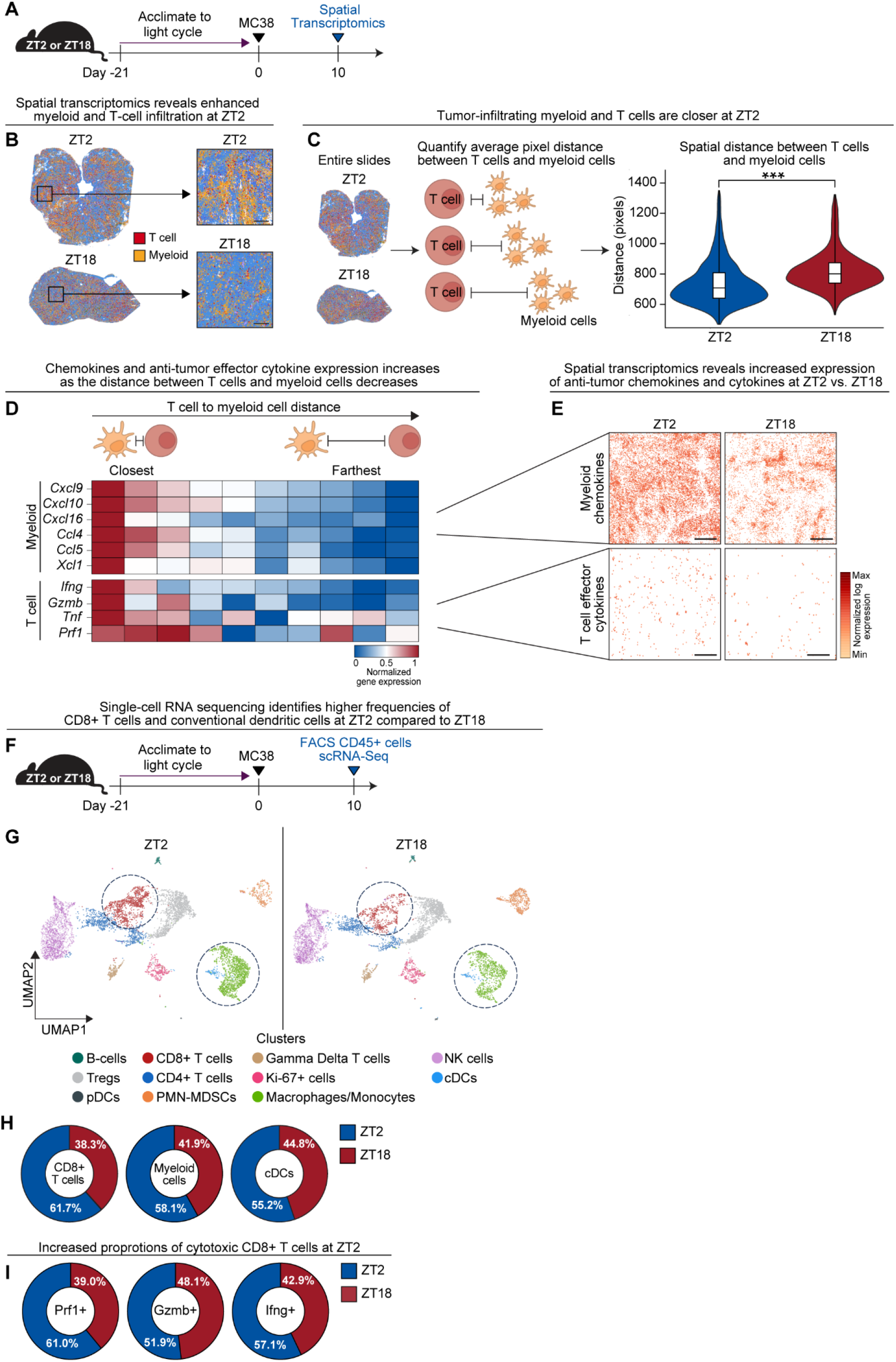
Time-of-day influences the spatial and cellular organization of the tumor immune microenvironment. **(A)** Schematic overview of the spatial transcriptomics experiment. MC38 tumors were inoculated into C57BL/6J mice (female, 8 weeks old) at ZT3 as in Fig. 1A. Tumors of equivalent size and weight were harvested on day 10 at ZT2 or ZT18 for spatial transcriptomics (Visium HD). **(B)** Spatial transcriptomic images with all annotated clusters shown. Entire tumor sections (low magnification, left) and higher-resolution images (high magnification, right), showing immune cell distributions at ZT2 and ZT18. **(C)** Experimental overview of the method to quantify spatial distances between T cells and neighboring myeloid cells at ZT2 and ZT18 (left and center). Violin plots with quantification of the distances between individual T cells and neighboring myeloid cells across entire tumor sections at ZT2 and ZT18 (right). **(D)** Heatmap examining gene expression of chemokines and effector cytokines in myeloid cell and T cell clusters, stratified into 10 distance quantiles based off spatial distances in (C). Each column represents a different distance quantile from closest together (far left column) to farthest apart (far right column). **(E)** Spatial feature plot showing average expression of chemokines in myeloid cells (*Cxcl9, cxcl10, cxcl16, ccl4, ccl5, xcl1)* and effector cytokines in T cells (*Ifng, gzmb, tnf, prf1*) in tumors collected at ZT2 vs. ZT18. **(F)** Schematic overview of single-cell RNA sequencing (scRNA-seq) of tumor-infiltrating CD45^+^ immune cells from tumors collected at ZT2 or ZT18. Tumors from 5 mice per time point were digested into single cell suspensions and pooled. CD45^+^ immune cells were isolated via FACS and sent for scRNA-sequencing. **(G)** UMAP plots of immune cell clusters from each time point; clusters annotated based on canonical gene expression (see Fig. S3). Dotted circles highlighting CD8^+^ T cell and myeloid cell clusters (cDC and monocyte/macrophage clusters). **(H)** Proportional contribution of CD8^+^ T cells, myeloid cells and cDCs at ZT2 versus ZT18. **(I)** Proportional contribution of effector cytokine (*Ifng, gzmb, prf1)* expressing CD8^+^ T cells at ZT2 versus ZT18. ZT, zeitgeber time; cDC, conventional dendritic cell; scRNA-seq, single cell RNA sequencing, UMAP, uniform manifold approximation and projection. Black scale bars represent 200 µm. Statistical analysis by Mann–Whitney test. **\*\*\***p < 0.001.

Spatial analysis of immune populations revealed that T cells were significantly closer to neighboring myeloid cells in ZT2 tumors than in ZT18 tumors (Fig. 2C), suggesting that time-of-day influences immune cell proximity and may promote increased immune cell crosstalk (*28, 29*).

Pathway analysis of differentially expressed genes showed that ZT2 tumors were enriched for immune-related transcriptional programs, including interferon-γ and chemokine signaling pathways (Fig. S2D), both of which are critical for T-cell activation and recruitment into the TIME. To link spatial proximity with functional consequences, we stratified gene expression by immune cell type and distance. Chemokine (CXCL9, CXCL10, CCL4, CCL5) and cytotoxic effector cytokine (interferon-γ, granzyme B, TNF-α, perforin) gene expression was highest in T cells and myeloid cells in close spatial proximity compared to immune cells that were farther apart (Fig. 2D). These same genes were significantly upregulated at ZT2 compared to ZT18 (Fig. 2E; Fig. S2E), consistent with enhanced immune activation during the inactive phase. Together, these findings suggest that differences in immune cell abundance, spatial organization, and activation state may underlie the improved ICT efficacy observed at ZT2.

To supplement these spatial transcriptomic findings, we performed single-cell RNA sequencing (scRNA-seq) on tumor-infiltrating CD45^+^ leukocytes isolated from tumors collected at ZT2 and ZT18 (Fig. 2F). Uniform Manifold Approximation and Projection (UMAP) analysis identified 11 transcriptionally distinct immune cell populations (Fig. 2G; Fig. S3A, B, C and D). ZT2 tumors were enriched for CD8^+^ T cells, monocytes/macrophages, and conventional dendritic cells (cDCs), whereas ZT18 tumors showed increased proportions of polymorphonuclear myeloid-derived suppressor cells (PMN-MDSCs) and plasmacytoid dendritic cells (Fig. 2G and H; Fig. S3E). Gene expression analysis confirmed increased chemokine expression in myeloid populations (Fig. S3F) and elevated frequencies of effector cytokine-producing CD8^+^ T cells at ZT2 (Fig. 2I).

In summary, these data suggest that time-of-day alters the cellular composition, spatial organization, and functional status of the TIME, providing a mechanistic basis for the enhanced ICT response observed during the inactive phase.

### Time-of-day-dependent dendritic cell and CD8^+^ T-cell infiltration shapes the TIME

We performed immune profiling (flow cytometry, immunohistochemistry (IHC), and multiplex cytokine analysis) on tumors collected at multiple timepoints to define protein-level differences in the TIME (Fig. 3A). To minimize confounding by tumor burden, only tumors with equivalent weights were analyzed (Fig. S4A). Flow cytometric profiling revealed significantly higher proportions (Fig. 3B and C) and absolute numbers (Fig. 3D) of live CD45^+^ tumor-infiltrating leukocytes (TILs) at ZT2 compared to ZT18, indicating marked time-of-day variation in overall immune infiltration.

**Figure 3.**
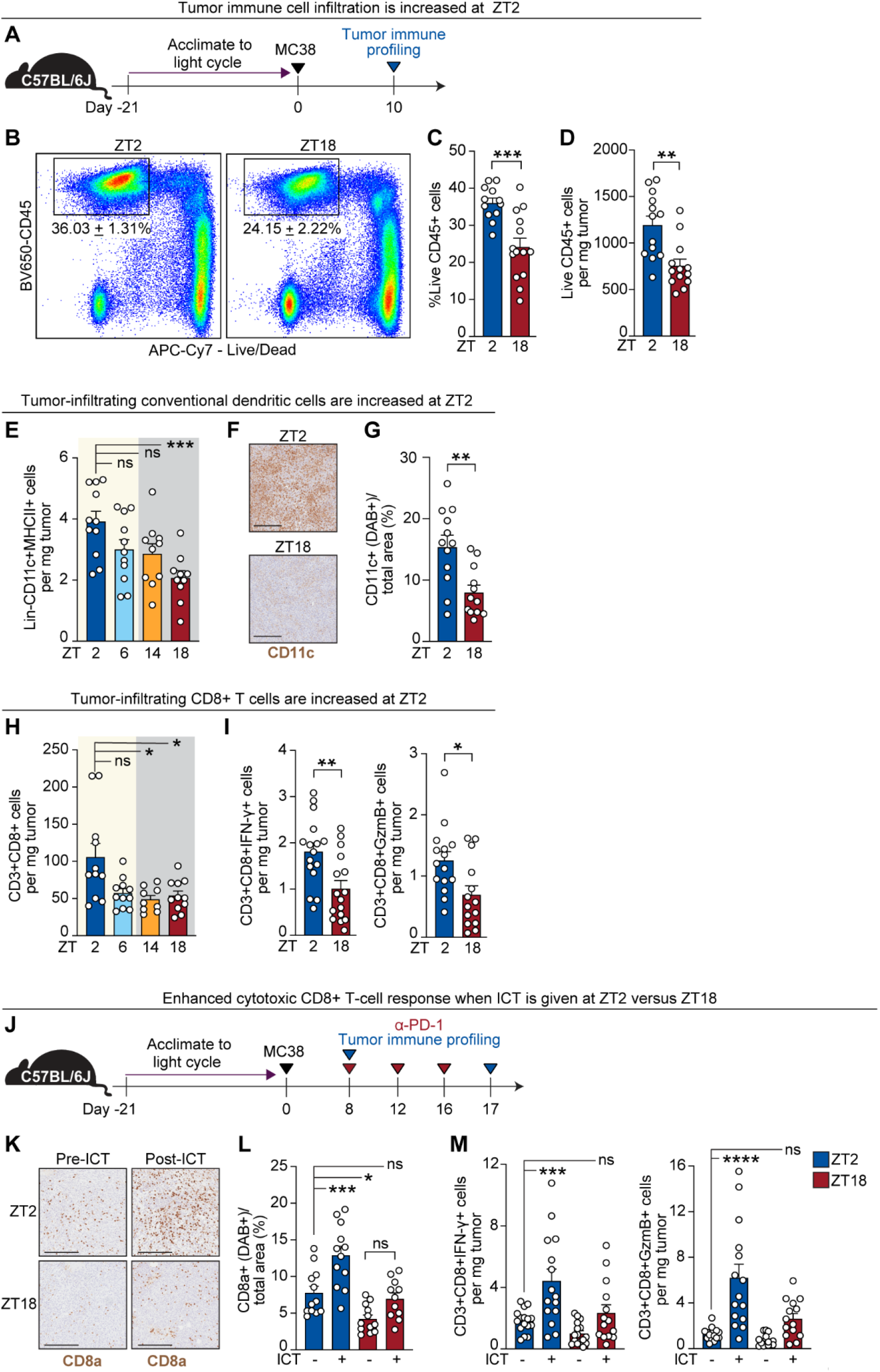
Time-of-day-dependent dendritic cell and CD8^+^T-cell infiltration shapes the TIME. **(A)** Schematic overview for tumor immune profiling. MC38 tumors were inoculated into C57BL/6J mice as in Fig. 1A. Tumors were harvested at indicated times of day (ZT2, ZT6, ZT14, ZT18) for immune profiling by flow cytometry, immunohistochemistry (IHC), or multiplex cytokine analysis. **(B)** Representative flow cytometry plots of live CD45^+^ tumor-infiltrating leukocytes (TILs) at ZT2 and ZT18. **(C)** Quantification of the frequency of CD45^+^ TILs at ZT2 versus ZT18; n = 12–14 mice per group. **(D)** Quantification of the absolute number of CD45^+^ TILs at ZT2 versus ZT18; n = 12–14 mice per group. **(E)** Quantification of tumor-infiltrating conventional dendritic cells (cDCs; Lineage?CD11c^+^MHCII^+^) across four time points (ZT2, ZT6, ZT14, ZT18) by flow cytometry; n = 10–11 mice per group. **(F)** Representative IHC images of CD11c^+^ staining in tumors harvested at ZT2 and ZT18. **(G)** Quantification of CD11c^+^ area (DAB^+^ signal) normalized to total tumor area; each point represents one quarter of a tumor section; n = 12 data points per group from 3 separate tumors. **(H)**Quantification of tumor-infiltrating CD8^+^ T cells (CD3^+^CD4^-^CD8^+^) across four time points (ZT2, ZT6, ZT14, and ZT18) by flow cytometry; n = 9–11 mice per group. **(I)** Quantification of IFN-γ^+^ (left) and granzyme B^+^ (right) CD8^+^ T cells in tumors at ZT2 versus ZT18; n = 14–16 mice per group. **(J)** Schematic overview of protocol used to assess CD8^+^ T cell dynamics in response to ICT administered at ZT2 or ZT18. **(K)** Representative IHC images of CD8a^+^-expressing cells before and after ICT at ZT2 or ZT18. **(L)** Quantification of CD8a^+^ area (DAB^+^ signal) normalized to total tumor area; each point represents one quarter of a tumor section; n = 11–12 data points per group from 3 separate tumors. **(M)** Quantification of IFN-γ^+^ (left) and granzyme B^+^ (right) CD8^+^ T cells in tumors before and after ICT at ZT2 or ZT18; n = 14–16 mice per group. ZT, zeitgeber time; ICT, immune checkpoint inhibitor therapy; IHC, immunohistochemistry; TIL, tumor-infiltrating leukocyte; cDC, conventional dendritic cell; DAB, 3’3’-diaminobenzidine. Statistical analysis by unpaired Student’s t test (C), Mann-Whitney (D, G, I, L) or one-way ANOVA (E, H, L, M). Points represent results from individual mice unless otherwise specified. Bars denote mean ± SEM. Black scale bars represent 200 µm. All data are representative of ≥2 independent experiments. *p < 0.05; **p < 0.005; ***p < 0.001; ****p < 0.0001. ns, not significant.

We next characterized myeloid populations across four timepoints (Fig. S4B for gating strategy), with a particular focus on CD11c^+^MHCII^+^ cDCs — key orchestrators of anti-tumor immunity and ICT response (*30, 31*). cDC abundance peaked at ZT2 and was significantly higher than at ZT18 (Fig. 3E). IHC confirmed greater DC infiltration at ZT2 via CD11c staining (Fig. 3F and G). No time-of-day differences were observed for macrophages or PMN-MDSCs, although Ly6C^+^ monocytes increased at ZT2 (Fig. S4C).

The shifts in myeloid composition were accompanied by functional changes. Chemokine concentrations in tumor lysates, assessed by Luminex-based multiplex analysis, were significantly elevated during the inactive phase (ZT2–ZT6) compared to the active phase (ZT14–ZT18) (Fig. S4D). This pattern is consistent with both increased myeloid abundance and enhanced functional output during the inactive period.

We then asked whether CD8^+^ T-cell infiltration and activation also varied by time of day, given their dependency on cDC priming and their central role in anti-tumor immunity (*6*). Indeed, CD8^+^ T cells were more abundant in tumors collected at ZT2, both in total numbers (Fig. 3H) and in the subset expressing effector cytokines, compared to ZT18 (Fig. 3I; see Fig. S5A for gating). Tumor lysates from ZT2 also contained higher concentrations of TNF-α, granzyme B, and IFN-γ (Fig. S5B). Additional lymphoid subsets (including CD4^+^ T cells, B cells, and NK cells) were similarly enriched at ZT2 relative to ZT18 (Fig. S5C).

Given that CD8^+^ T-cell recruitment, priming, and effector function are critical determinants of ICT efficacy, we hypothesized that time-of-day differences in baseline CD8^+^ T-cell infiltration might dictate ICT response. To test this, we treated MC38-bearing mice with ICT at ZT2 or ZT18 and collected tumors after three doses (Fig. 3J). IHC of untreated tumors confirmed higher baseline CD8a^+^ T cell density at ZT2 versus ZT18 (Fig. 3K left images). ICT given at ZT2 induced a robust expansion of intratumoral CD8a^+^ T cells, whereas ICT at ZT18 failed to elicit a comparable increase (Fig. 3K and L). Moreover, ZT2 treatment significantly enhanced the frequency of IFN-γ^+^ and granzyme B^+^ CD8^+^ T cells (Fig. 3M; Fig. S5D), indicating a stronger cytotoxic CD8^+^ T-cell response.

Together, these findings demonstrate that cDC and CD8^+^ T cell abundance in the TIME is temporally regulated and may underlie the enhanced ICT response observed during the inactive phase.

### Dendritic cell and CD8^+^ T-cell circadian clocks are required for ICT efficacy

The pronounced time-of-day-dependent differences in dendritic cell and CD8^+^ T-cell function (Figs. 2 and 3) prompted us to ask whether these were driven by immune cell–intrinsic circadian clocks. BMAL1 is a core circadian transcription factor; its deletion abolishes rhythmicity (*11*). To test whether ICT efficacy depends on immune cell-intrinsic clocks, we generated conditional knockout mice lacking *Bmal1* in DCs (*Bmal1*^*flox/flox*^ × CD11c-Cre, *Bmal1*^*ΔDC*^ mice) or CD8^+^ T cells (*Bmal1*^*flox/flox*^ × CD8a-Cre, *Bmal1*^*ΔCD8*^ mice). *Bmal1* deficiency in DCs and CD8^+^ T cells has been shown to impair immune cell migration, antigen presentation, cytokine/chemokine production, and responses to pathogens, tumor engraftment, and vaccination (*12–16, 32, 33*).

We first assessed the role of the DC circadian clock in regulating ICT response (Fig. 4A). ICT was less effective in *Bmal1*^*ΔDC*^ mice, with increased tumor growth (Fig. 4B) and decreased survival (Fig. 4C), compared to *Bmal1*^*fl/fl*^ controls treated at ZT2. Tumor immune cell profiling in *Bmal1*^*ΔDC*^ mice confirmed a blunted cytotoxic CD8^+^ T-cell response, with fewer IFN-γ^+^ and Granzyme B^+^ CD8^+^ T cells after ICT (Fig. 4D). In contrast, *Bmal1*^*fl/fl*^ controls retained time-of-day differences in ICT efficacy, mirroring wild-type mice results (Fig. 1B, C and D). Notably, this variation was abrogated in *Bmal1*^*ΔDC*^ mice, indicating that DC clocks are required for the time-of-day differences in ICT response.

**Figure 4.**
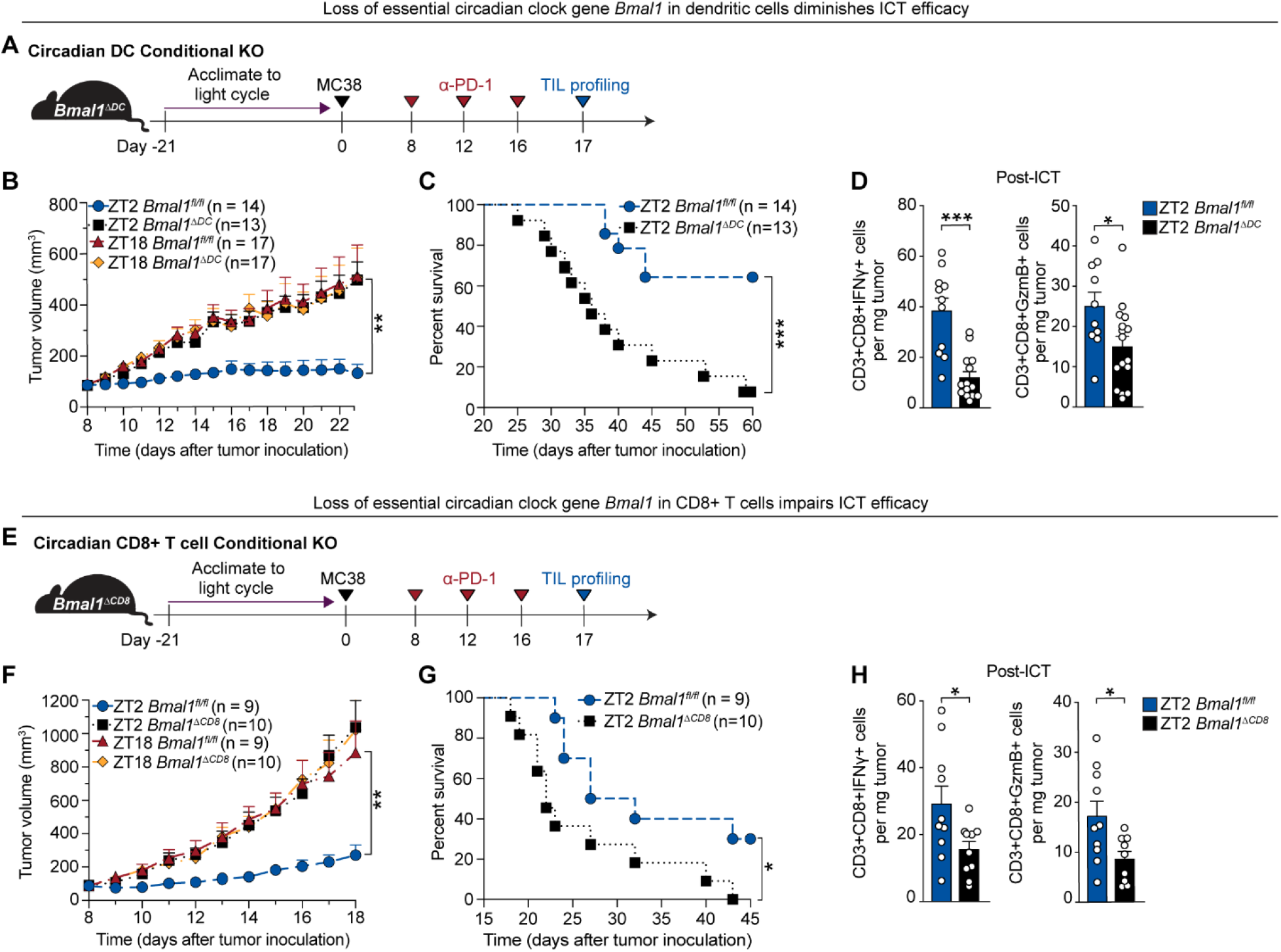
Dendritic cell and CD8^+^ T-cell circadian clocks are required for ICT efficacy. **(A)** Schematic overview of protocol to assess ICT (anti–PD-1) response in MC38 tumor-bearing mice with dendritic cell–specific deletion of the core clock gene *Bmal1* (*Bmal1*^*fl/fl*^ x CD11c-cre mice, *Bmal1*^*ΔDC*^). **(B)** Tumor growth of *Bmal1*^*ΔDC*^ and *Bmal1*^*fl/fl*^ control mice treated with ICT at ZT2 or ZT18; n = 13–17 mice per group. **(C)** Survival of *Bmal1*^*ΔDC*^ and *Bmal1*^*fl/fl*^ control mice treated with ICT at ZT2; n = 13–17 mice per group. **(D)** Quantification of IFN-γ^+^ (left) and granzyme B^+^ (right) CD8^+^ T cells in tumors of *Bmal1*^*ΔDC*^ or *Bmal1*^*fl/fl*^ mice after ICT at ZT2 via flow cytometry; n = 10–16 mice per group. **(E)** Schematic overview of protocol to assess ICT (anti–PD-1) response in MC38 tumor-bearing mice with CD8^+^ T cell–specific *Bmal1* deficiency (*Bmal1*^*fl/fl*^ x CD8a-cre mice, *Bmal1*^*ΔCD8*^). **(F)** Tumor growth of *Bmal1*^*ΔCD8*^ vs. *Bmal1*^*fl/fl*^ mice treated with ICT at ZT2 or ZT18; n = 9–10 mice per group. **(G)** Survival of *Bmal1*^*ΔCD8*^ and *Bmal1*^*fl/fl*^ mice treated with ICT at ZT2; n = 9–10 mice per group. **(H)** Frequency of IFN-γ^+^ (left) and granzyme B^+^ (right) CD8^+^ T cells in tumors of *Bmal1*^*ΔCD8*^ or *Bmal1*^*fl/fl*^ mice after ICT; n = 9–11 mice per group. KO, knock out; ZT, zeitgeber time; TIL, tumor-infiltrating leukocyte; ICT, immune checkpoint inhibitor therapy; Statistical analysis by Mann–Whitney (B, D, F, H) or log-rank test (C, G). Points represent results from individual mice. Bars denote mean ± SEM. Data are representative of ≥2 independent experiments. *p < 0.05; **p < 0.01 ***p < 0.001.

We next tested whether the CD8^+^ T-cell clock similarly regulates ICT efficacy (Fig. 4E). *Bmal1*^*ΔCD8*^ mice showed impaired ICT efficacy, with greater tumor burden (Fig. 4F) and reduced survival (Fig. 4G) compared to *Bmal1*^*fl/fl*^ controls at ZT2. Additionally, *Bmal1*^*ΔCD8*^ mice failed to show time-of-day differences in ICT efficacy. CD8^+^ T cell profiling revealed reduced numbers of IFN-γ^+^ and granzyme B^+^ CD8^+^ T cells in *Bmal1*^*ΔCD8*^ mice compared to controls after ICT (Fig. 4H). Together, these findings demonstrate that circadian clocks in DCs and CD8^+^ T cells are essential for ICT efficacy and its time-of-day dependence.

### Circadian regulation of CX3CL1–CX3CR1 signaling modulates time-of-day differences in tumor immune infiltration and ICT efficacy

As DC clocks were required for ICT efficacy and are essential for mediating CD8^+^ T cell recruitment, we asked whether the DC-intrinsic clock regulates specific chemokines that are known to shape TIME composition and could regulate time-of-day differences in ICT efficacy. We performed RNA-sequencing of sorted tumor-infiltrating dendritic cells (CD45^+^Lineage^−^CD11c^+^) collected at ZT2 and ZT18 in wild-type mice (Fig. 5A). Kyoto Encyclopedia of Genes and Genomes (KEGG) pathway analysis revealed enrichment of circadian entrainment and chemokine signaling pathways at ZT2 (Fig. S6A). Among differentially expressed genes, *Cx3cl1* (fractalkine) was the only chemokine significantly upregulated at ZT2 (Fig. 5B) – notable given its known role in immune cell trafficking (*29, 34*). CX3CL1 protein levels were also elevated in tumor lysates at ZT2 compared to ZT18 (Fig. 5C).

**Figure 5.**
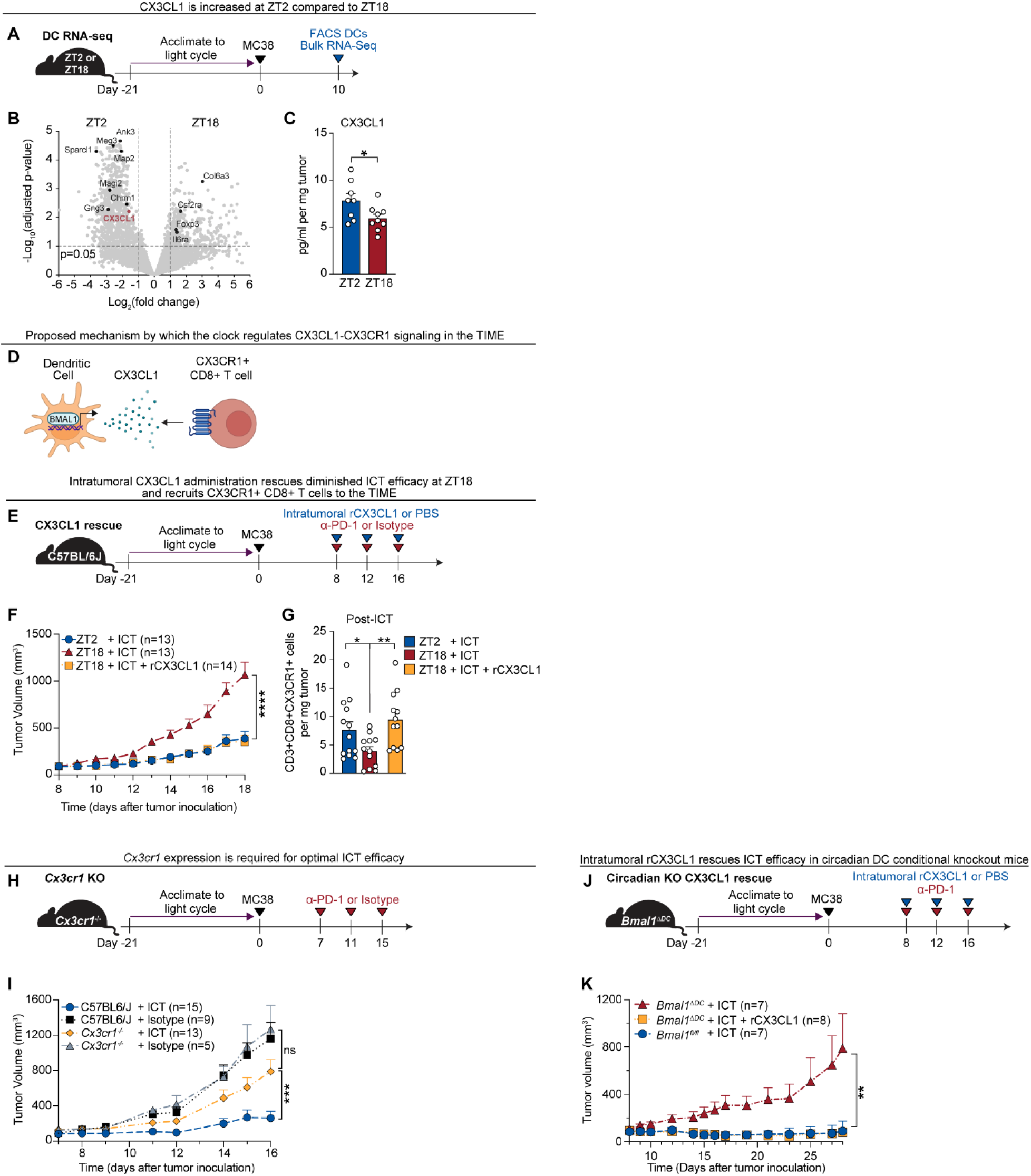
Circadian regulation of CX3CL1–CX3CR1 signaling modulates time-of-day differences in tumor immune infiltration and ICT efficacy. **(A)** Schematic overview of protocol to assess transcriptional changes in tumor-infiltrating DCs at ZT2 and ZT18. C57BL/6J mice were implanted with MC38 tumors and on day 10, tumor-infiltrating DCs (CD45^+^Lineage^-^CD11c^+^ cells) were isolated using FACS and sent for RNA sequencing. **(B)** Volcano plot visualizing the changes in gene expression between tumor-infiltrating dendritic cells at ZT2 vs. ZT18. *Cx3cl1* is highlighted in red. **(C)** Quantification of CX3CL1 protein levels in tumors harvested at ZT2 and ZT18, measured by Luminex assay; n = 8–9 mice per group. **(D)** Proposed mechanism by which the circadian clock regulates *Cx3cl1* expression in DCs to recruit CX3CR1^+^ CD8^+^ T cells to tumors. **(E)** Schematic overview of the protocol used to test the impact of intratumoral rCX3CL1 on ICT response and CX3CR1^+^CD8^+^ T cell recruitment. C57BL6/J mice were implanted with MC38 tumors and were injected intraperitoneally with 200 µg anti–PD-1 or isotype ± 100ng of intratumoral rCX3CL1 or PBS vehicle control at ZT2 or ZT18 on days 8, 12 and 16. Tumors were collected for flow cytometry after ICT and rCX3CL1 treatment on day 17. **(F)** Tumor volumes of tumor-bearing mice from (E); n = 13–14 mice per group. See Fig. S6E for all groups. **(G)** Quantification of tumor-infiltrating CX3CR1^+^CD8^+^ T cells following ICT ± rCX3CL1 at ZT2 or ZT18; n = 12–13 mice per group. **(H)** Schematic overview of the protocol to assess the impact of *Cx3cr1* deficiency on ICT efficacy. *Cx3cr1* knockout (*Cx3cr1*^-/-^) and C57BL/6J control mice were given MC38 tumors and treated at ZT2 with 200 µg anti–PD-1 or isotype control on days 7, 11, and 15. **(I)** Tumor volumes of tumor-bearing mice from (H); n = 5–15 mice per group. **(J)** Schematic overview of the protocol to assess the impact of rCX3CL1 on ICT efficacy in dendritic cell-specific circadian knockout mice (*Bmal1*^*ΔDC*^) versus control mice (*Bmal1*^*fl/fl*^). ICT was administered at ZT2 in all groups. **(K)**Tumor volumes of tumor-bearing mice from (J); n = 7–8 per group. ZT, zeitgeber time; FACS, fluorescence activated cell sorting; DC, dendritic cell; RNA-seq, RNA sequencing; ICT, immune checkpoint inhibitor therapy; TIME, tumor immune microenvironment; rCX3CL1, recombinant CX3CL1; KO, knockout. Statistical analysis by Student’s *t* test (C), or Mann-Whitney test (F, G, H, J, K). Points represent individual mice. Bars denote mean ± SEM. All data are representative of ≥2 independent experiments. *p < 0.05; **p < 0.01; ***p < 0.001; ****p < 0.0001. ns, not significant.

CX3CL1 recruits immune cells (including CD8^+^ T cells, NK cells, and DCs) through binding its sole receptor CX3CR1 (*29, 34*). Although CX3CL1 has been linked to anti-tumor immunity (*35*), its role in ICT response is undefined. Promoter analysis of *Cx3cl1* revealed six putative BMAL1-binding E-box motifs (Fig. S6B) (*11, 12*), suggesting direct circadian regulation. ChIP-qPCR confirmed BMAL1 binding to the *Cx3cl1* promoter in DCs (Fig. S6C), identifying *Cx3cl1* as a transcriptional target of the DC clock.

We next tested whether CX3CL1 regulates time-of-day differences in ICT efficacy (Fig. 5E). Intratumoral recombinant CX3CL1 (rCX3CL1) administered with ICT at ZT18 reduced tumor volume (Fig. 5F) and improved survival (Fig. S6D) to levels comparable to ZT2 ICT treatment. In contrast, rCX3CL1 alone had no effect on tumor growth in the absence of ICT (Fig. S6E).

We then asked whether CX3CL1 acts via CX3CR1^+^ CD8^+^ T cells, a putative biomarker of durable ICT responses in patients (*36-38*). scRNA-seq revealed higher *Cx3cr1* expression and a greater proportion of *Cx3cr1*-expressing cells within the CD8^+^ T cell cluster at ZT2 (Fig. S6F). Flow cytometry confirmed that intratumoral CX3CR1^+^ CD8^+^ T cell abundance peaked at ZT2 (Fig. S6G and H). Importantly, rCX3CL1 treatment at ZT18 restored CX3CR1^+^ CD8^+^ T cells to ZT2 levels (Fig. 5G).

To determine whether CX3CR1 is required for ICT efficacy, we treated *Cx3cr1*-deficient (*Cx3cr1*^-/-^) mice with ICT at ZT2 (Fig. 5H). Compared to wild-type controls, *Cx3cr1*^-/-^ mice exhibited impaired ICT responses and accelerated tumor growth (Fig. 5I). Finally, we asked whether rCX3CL1 could rescue dendritic cell clock deficiency. Strikingly, rCX3CL1 fully restored ICT responsiveness in *Bmal1*^*ΔDC*^ mice (Fig. 5J and K).

Together, these findings identify *Cx3cl1* as a direct transcriptional target of the DC clock and demonstrate that time-of-day-dependent differences in CX3CL1–CX3CR1 signaling regulate CD8^+^ T-cell recruitment to the TIME, driving temporal variation in ICT efficacy (Fig. S7).

## Discussion

This study demonstrates that the circadian clock regulates ICT efficacy by shaping the tumor immune microenvironment. We identify *Cx3cl1* as a direct transcriptional target of the DC clock and show that time-of-day differences in CX3CL1-CX3CR1 signaling, direct CD8^+^ T-cell recruitment into tumors. Administering ICT during the inactive phase (ZT2), when immune activity peaks, enhances immune infiltration, CD8^+^ T-cell effector function, and therapeutic efficacy (Fig. S7).

These findings unify and extend prior work implicating circadian regulation of anti-tumor immunity (*16, 39, 40*). *Fortin et al*. showed that circadian disruption drives intratumoral accumulation of PD-L1–expressing myeloid-derived suppressor cells (MDSC) and synchronization of anti–PD-L1 therapy with the peak of PD-L1-expresing MDSC infiltration enhances efficacy (*39*). Additionally, *Wang et al*. demonstrated circadian regulation of endothelial cell and T-cell clocks which coordinate rhythmic T-cell infiltration, leading to time-of-day differences in CAR-T and ICT responses (*40*). Our work builds on these observations by identifying a chemokine pathway—CX3CL1–CX3CR1—as a direct transcriptional output of the DC clock. This axis coordinates time-of-day-dependent CD8^+^ T-cell recruitment and modulates anti-tumor immunity in a manner that directly impacts ICT efficacy. Using multiple tumor models, immune cell–specific knockouts, and spatial and single-cell transcriptomics, we provide a mechanistic framework defining a pathway from immune cell–intrinsic circadian programs to ICT efficacy.

These mechanisms remain to be validated in human cancer. Although our study was limited to mouse models, the molecular circadian clock and the CX3CL1–CX3CR1 axis are conserved in humans. Longitudinal profiling of tumor-infiltrating or peripheral immune cells could identify circadian biomarkers predictive of response. In parallel, functional studies across additional tumor types and metastatic sites are needed to determine whether the CX3CL1–CX3CR1 pathway broadly influences TIME dynamics.

Clinical studies already suggest that treatment timing may impact immunotherapy outcomes (*18–20*). Retrospective analyses have reported improved survival in patients receiving ICT earlier in the day. A recent randomized clinical trial showed that non–small cell lung cancer patients treated with morning chemoimmunotherapy had significantly longer progression-free survival than those treated in the afternoon (*41*). These clinical findings align with our preclinical data showing enhanced ICT efficacy when administered early in the light phase (Fig. 1). Our findings provide a mechanistic foundation for these clinical observations and support the integration of treatment timing as a critical variable in future immunotherapy trials.

In summary, we identify the circadian clock as an actionable regulator of cancer immunotherapy. Aligning treatment with endogenous immune rhythms or targeting downstream effectors such as CX3CL1 may enhance ICT efficacy and overcome therapeutic resistance. These findings provide a mechanistic rationale for chrono-immunotherapy and support the incorporation of treatment timing into future clinical trials to optimize immunotherapy outcomes in cancer patients.

## Acknowledgments

We thank Prithvi Raj, Chiaoying Liang, Yang Liu and Indu Ramen from the UT Southwestern Genomics Core for assistance with spatial transcriptomics and RNA sequencing experiments. We thank the UT Southwestern Microarry Core for assistance with Luminex assays and the UT Southwestern McDermott Next Generation Sequencing Core for assistance with scRNA-sequencing. We used OpenAI’s ChatGPT (version 4.0 and 5.0) solely for language editing (e.g., grammar, clarity, and brevity) and for suggesting alternative phrasings during manuscript preparation. Components of some figures were made using BioRender.

## Author contributions

JNL, LVH, and AYK designed the research. JNL, TS, CD, JIR, NP, HW, WL, NP, AM, BH, PS, SP, JMS, and XVL performed the research. BME performed blinded histology image analysis. JST provided experimental materials. JNL, RW, JK, and XZ analyzed the data. JNL, CMZ, and AYK wrote the paper. All authors revised the manuscript and approved its final version.

## Data and materials availability

Spatial transcriptomics, single cell RNA sequencing and RNA sequencing data will be available from the Gene Expression Omnibus (GEO) repository (deposition of data currently underway). All other data are available in the main text or the supplementary materials.

## Supplementary Materials

Materials and Methods Figs. S1 to S7 Table S1

References (*42-55*)

