## Supplemental Data 1 for "Timing of immune checkpoint blockade shapes anti-tumor immunity via a clock-dependent chemokine axis"

**The PDF file includes:**

Materials and Methods
Supplementary Text
Figures S1 to S7
Table S1
References (42-55)

### Materials and Methods

#### Mice

C57BL/6J wild-type, *Bmal1*$^{fl/fl}$ (*42*), CD11c-cre (*43*), CD8a-cre (*44*), *Cx3cr1*$^{-/-}$ (*45*) mice were obtained from Jackson Laboratories and bred and maintained in the animal facility at the University of Texas Southwestern Medical Center. Transgenic mice (*Bmal1*$^{\Delta DC}$ (*46*) and *Bmal1*$^{\Delta CD8}$ (*47*)) were maintained as homozygous for *Bmal1*$^{flox/flox}$ and heterozygous for the applicable Cre. All mice were genotyped using the relevant PCR protocols from Jackson Laboratory prior to each experiment. All animals were kept on a 12-hour light-dark cycle and were fed standard mouse chow (Teklad 2916, irradiated) and water *ad libitum*. Sex-matched, 6–12-week-old mice were used for all experiments. To investigate multiple time points simultaneously, mice were housed in light-tight animal cabinets (ActiMetrics). Mice were housed for 21 days prior to the start of an experiment to acclimate to the light cycle. Treatment times correspond to Zeitgeber time (ZT), which refers to the timing relative to the light cycle in the animal facility (e.g. ZT2 refers to 2 hours after lights on and ZT18 refers to 6 hours after lights off). During the dark phase, all animal handling and experimentation (e.g. ICT administration) was done under a dim red light. Experiments were performed using protocols approved by the Institutional Animal Care and Use Committees of the UT Southwestern Medical Center

#### Cell Lines

B16-F10 cells (ATCC CRL-6475; RRID:CVC_0159), MC38 cells (gift from Dr. Todd Aguilera, University of Texas Southwestern Medical Center, RRID:CVCL_B288), and MC38-luciferase cells (ALSTEM Bio, RRID:CVCL_C8VZ) were grown at 37°C under 5% CO2 in DMEM medium supplemented with 10% heat-inactivated FBS (Gibco), 100 units/ml penicillin, and 100µg/ml streptomycin sulfate (Thermo Fisher). All cell lines routinely tested negative for *Mycoplasma* (ATCC PCR Mycoplasma Detection Kit). Only cells with a passage number <10 were used for experiments.

#### Preclinical tumor models and immune checkpoint inhibitor therapy

For the heterotopic MC38 model, $5 \times 10^5$ MC38 cells were inoculated subcutaneously into the right flank of mice. On day eight after tumor inoculation, mice were separated into treatment groups with equivalent tumor volumes ($100mm^3 \pm 30mm^3$). Mice not meeting tumor size criteria were excluded from the experiment. Mice were then injected with 200 µg anti-PD-1 antibody (RMP1-14, CD270, BioXcell) or isotype control (Rat IgG2a, BioXcell) intraperitoneally at indicated time points (ZTs). An additional two doses of ICT or isotype were given at four-day intervals.

For the orthotopic melanoma model, $2 \times 10^5$ B16-F10 cells were implanted subcutaneously into the right flank of mice. On day ten after tumor inoculation, mice were separated into treatment groups with equivalent tumor volumes ($100mm^3 \pm 30mm^3$). Mice not meeting tumor size criteria were excluded from the experiment. Mice were then injected with 200 µg anti-PD-1 antibody (RMP1-14, CD279, BioXcell) and 200 µg anti-CTLA-4 antibody (9D9, CD152, BioXcell) or isotype control (Rat IgG2a, κ or Mouse IgG2b, respectively, BioXcell) intraperitoneally at indicated time points (ZTs). An additional two doses of ICT or isotype were given at four-day intervals.

For both subcutaneous tumor models, tumor volumes were monitored every 1-2 days using a caliper. Tumor volume was calculated using measurements from a digital caliper and the following formula: (Length x width x width)/2. Loss of survival was defined as death (with moribund mice being euthanized) or when tumor diameter >2 cm in any dimension or total tumor volume >1800 $mm^3$.

For the orthotopic rectal MC38-Luc model, 1 x $10^6$ MC38-Luc cells (luciferase positive MC38 cells) were resuspended in Cultrex (R&D systems) and serum-free DMEM at a 1:1 ratio. Mice were then anesthetized with vaporized isoflurane and the lumen of the mouse rectum was exposed (without having to make a surgical incision) by pushing on the pelvis of the mouse and using a sterile pipette tip. Using a 27g needle, 30 µl of the cell mixture was injected into lumen of the mouse rectum. Seven days after tumor inoculation, the mice were prepared for bioluminescent imaging. They were first injected intraperitoneally with 200 µl of D-luciferin (15mg/ml, GOLD Bio) dissolved in sterile PBS. They were then anesthetized with vaporized isoflurane and eight minutes later, bioluminescent mouse images were captured using a bioluminescent imager (IVIS Spectrum, Perkin Elmer). The bioluminescent signal (average radiance) of each rectal tumor was quantified using the LivingImage™ software (Perkin Elmer, see Fig. S1C for representative mouse images). Mice were separated into different treatment groups with equivalent average bioluminescence per group so that they each started with equivalent tumor burden. On day 8 after tumor inoculation, mice were then injected with 200 µg anti-PD-1 antibody (RMP1-14, CD270, BioXcell) or isotype control (Rat IgG2a, BioXcell) intraperitoneally at indicated time points (ZTs). An additional two doses of ICT or Isotype were given at four-day intervals on day 12 and day 16 post tumor inoculation. The mice underwent further bioluminescent imaging sessions on day 11, day 15 and 19 post tumor inoculation to monitor response to ICT. On day 19, the mice were euthanized, and rectal tumors were harvested to quantify tumor weight.

**Tumor-infiltrating leukocyte isolation**

Single cells from dissected tumor tissue were isolated using the gentleMACS™ Octo Dissociator with Heaters and mouse tumor dissociation kit (Miltenyi Biotec), according to the manufacturer's protocol. Briefly, tumor-bearing mice were euthanized ($CO_2$ inhalation) at indicated time points. The entire tumor was removed, and the tumor draining lymph node and excess fat were excluded. being careful to remove the tumor draining lymph node and any excess fat. The tumor was then weighed and subsequently cut into 2–4 mm pieces. The pieces of tumor were mechanically and enzymatically dissociated using enzymes from the tumor dissociation kit and placed on the gentleMACS™ Octo Dissociator with Heaters (37_m_TDK_1 protocol) for 40 minutes. The dissociated tumor was then passed through a 70 µm MACS Smart Strainer (Miltenyi Biotec) into a 15 ml conical tube and washed with 10 ml of RPMI medium with 10% FBS. The cell suspension was centrifuged at 400 x g, 4°C for 10 min. The supernatant was carefully aspirated, and the cell pellet was suspended in RBC lysis buffer (Invitrogen) and incubated at room temperature (RT) for 2 min to remove erythrocytes. The cells were washed with 10 ml of PBS and centrifuged at 400 x g, 4°C for 5 min. Cell pellets were suspended in a 40% Percoll solution to remove fat and necrotic tissue. Suspensions were centrifuged at 350 x g at RT for 20 min (ascending rate: 5; descending rate: 0). The supernatant was then carefully aspirated, and the cell pellet was washed once with PBS prior to proceeding with further experiments (e.g. flow cytometry).

**Flow cytometry analysis**

Single-cell suspensions of cells were transferred to V-bottom 96 well plates (Corning) and centrifuged at 400 x g, 4°C for 10 min. For surface staining, cells were incubated with Ghost 780 live/dead dye (Tonbo Biosciences) diluted in PBS (1:1000) at RT for 15 minutes. After washing with PBS, cells were incubated with anti-CD16/CD32 (Fcγ receptor) blocking antibody (Biolegend), diluted 1:500 in FACS buffer (PBS supplemented with 3% heat-inactivated fetal bovine serum (FBS) and 1mM EDTA) for 10 min at 4℃ in the dark followed by surface staining with primary antibodies (1:200) for 30 min at room temperature in the dark. For surface staining, 10 µl of Horizon Brilliant Stain Buffer Plus (BD Biosciences) was added to each well to minimize staining artifacts. After staining, cells were then washed with FACS buffer twice and suspended in FACS buffer prior to flow cytometry.

For intracellular staining, single cell suspensions were transferred to U-Bottom 96 well plates (Corning) and incubated in 200µl RPMI supplemented with 10% FBS, 1x Cell Activation Cocktail (Biolegend, cocktail contains Phorbol 12-Myristate 13-Acetate (PMA), Ionomycin and Brefeldin-A) at 37°C with 5% CO2 for 4.5 hours. After incubation for 4.5 hours, cells were washed with PBS. Cells were then incubated with Ghost Dye Red 780 live/dead (Tonbo Biosciences) diluted in PBS (1:1000) at RT for 15 minutes. After washing with FACS buffer, cells were incubated with anti-CD16/CD32 (Fcγ receptor) blocking antibody, diluted in FACS buffer for 10 min at 4℃ in the dark followed by surface staining (as above) for 30 min at room temperature in the dark. Cells were then fixed and permeabilized using the Cyto-fast Fix-Perm Buffer Set (Biolegend) according to the manufacturer's instructions. Cells were then stained with antibodies targeting intracellular proteins (1:100) for 30 min at room temperature. Cells were washed with FACS buffer twice and suspended in FACS buffer prior to flow cytometry. All flow cytometry experiments were performed on a Novocyte Advanteon (Agilent Technologies). Flow cytometry gating strategy for the various experiments used in this study are detailed in Fig. S4B, S5A, and S6H. Antibodies used for flow cytometry are listed in Table S1.

### Single cell RNA sequencing of tumor-infiltrating immune cells

Tumors were harvested, weighed and single cell suspensions were prepared as above. Prior to single cell RNA sequencing, tumor-infiltrating immune cells ($CD45^+$ cells) were isolated using fluorescence activated cell sorting (FACS) using a FACSymphony S6 cell sorter (BD Biosciences). In order to get representative samples across multiple mice, single cell suspensions from five mouse tumors of equivalent weight were collected at ZT2 and ZT18 and pooled prior to sorting. Live $CD45^+$ cells were sorted into 1% BSA in PBS. Purified cellular suspensions of $CD45^+$ cells were loaded onto a 10x Genomics Chromium Instrument at concentration of 1200 cells/µl. Single cell RNA-seq libraries were prepared using the Chromium Single Cell 3' v3.1 Gene Expression Kit (10x Genomics) according to manufacturer's instructions. Single-cell RNA-seq libraries were sequenced on an Illumina NovaSeq 6000 using paired end reads.

### Single cell RNA sequencing clustering and analysis

Cell Ranger Software (v6.0) was used for sample demultiplexing, barcode processing, and single cell counting. Cell Ranger count was used to align samples to the reference mouse genome (UCSC mm10). The Seurat (v5.1.0) package in R (v4.3.1) was used for downstream analysis including clustering. Cells with mitochondrial content greater than 20% were removed from further analysis. In addition, cells with low Unique Molecular Identifier (UMI) (<6000) and gene number per cell (<100) were also excluded from further analysis. After filtering and integration (*48*), the dataset shown in the Figure 2 or Figure S3 contained a total of 12,790 cells (for ZT2 and for ZT18) with a median UMI of 5,819 and median of 1,784 genes per cell. Uniform Manifold Approximation and Projection (UMAPs) were computed in Seurat using 30 dimensions. Cluster marker genes were identified using the FindAllMarkers function in Seurat, employing Wilcoxon rank-sum tests. Immune cell clusters were annotated based off canonical marker gene expression (See Fig. S3). Visualization of scatterplots were implemented using custom R scripts.

**Histology and Immunohistochemistry**

Subcutaneous MC38 tumors from C57BL/6J mice were collected at indicated ZTs prior to ICT initiation or 24 hours after the third dose of ICT (200 µg anti-PD-1) and carefully placed into histology cassettes. Histology cassettes were immediately immersed in 4% paraformaldehyde in PBS and placed on a laboratory rocker at room temperature for 48 hours. Samples were transferred to 70% ethanol in water. The fixed tumor tissue was dehydrated, cleared, and infiltrated with paraffin. Samples were embedded for maximum surface area and sectioned longitudinally. A battery of three slides was prepared from each block. 5 µm serial sections were collected for routine hematoxylin and eosin (H&E), and CD8a and CD11c immunohistochemistry to characterize the presence of tumor-infiltrating $CD8a^+$ T cells or $CD11c^+$ dendritic cells. H&E regressive staining was performed using Leica-Surgipath Selectech reagents (Hematoxylin 560, Define Concentrate, Blue Buffer, Alcoholic Eosin Y 515, Deer Park, IL) on a Sakura DRS601 x-y-z robotic stainer. Slides for CD8a and CD11c were processed separately due to same-species-primary-antibody challenges and differing antigen retrieval methods. IHC slides were deparaffinized, and antigens retrieved by heating in pH 9.0, 10mM Tris/1mM EDTA (CD8a) or pH 7.4, 0.05% Citraconic Anhydride (CD11c), and sections were blocked against secondary antibody host-serum affinity utilizing commercially available blocking reagents (Impress Block Normal Goat Serum, Vector Laboratories, Burlingame CA). Following blocking, sections were subjected to either CD8a primary antibody (1:200, rabbit monoclonal EPR21769, Abcam Inc., Waltham, MA) or CD11c antibody (1:50, rabbit monoclonal D1V9Y, Cell Signaling Technology, Danvers, MA) and incubated overnight at 4°C. Subsequent biotin/streptavidin-peroxidase detection of bound primaries (CD8a or CD11c) were conducted the following day according to manufacturer's instructions (Immpress Excel Amplified Polymer Staining Kit, Anti-Rabbit IgG, Peroxidase, Vector Laboratories, MP-7601). At the conclusion of IHC staining, nuclei were lightly counterstained with hematoxylin (1:12 dilution of Leica Selectech Hematoxylin 560) and blued in running tap water. Slides were thereafter dehydrated, cleared, and mounted with coverslips using synthetic mounting media.

**Light Microscopy and IHC Quantification**

H&E, CD8a, and CD11c IHC slides were imaged using an Aperio CS2 slide scanner (Leica Biosystems, Deer Park IL, USA) at 40x magnification. Images were obtained at 10x and 20x using ImageScope software version 12.3 (Leica Biosystems Deer Park Illinois, USA). To qualitatively confirm time-of-day differences in $CD8a^+$ T cells or $CD11c^+$ dendritic cells, images were independently reviewed by pathologist Bret M. Evers M.D. Ph.D., UTSW Histopathology Core, who was blinded to the treatment groups. For quantification of $CD8a^+$ or $CD11c^+$ staining, CD8a and CD11c stained tumor sections were reviewed and imaged by a blinded observer using bright-field illumination on a PrimeHisto XE scanner (Pacific Image Electronics, Torrance, CA). Individual RGB images were saved as 8-bit TIFF and were collected at 5000 DPI using HistoView software version 1.00.75. Resulting images were uniformly contrast and gamma adjusted in Adobe Photoshop 25.12.1 to improve tissue versus non-tissue contrast before import into Fiji ImageJ2 (version 2.14.0/1.54f). Within Fiji ImageJ2, RGB image information was converted to grayscale using RGB stack function, and blue-channel information was chosen for offering the best DAB-contrast dynamic-range. Respective DAB-immunostained areas were highlighted by setting threshold for inclusion intensity values between 0 (absolute black) and 163 (medium gray) and the morphometric area recorded in pixels. Working with the same image, total tissue morphometric area was similarly highlighted but by setting threshold inclusion intensity values between 0 and 249 (lighter gray) and recorded in pixels. From each microscopy field, CD8a positive area versus total tissue area (or CD11c positive area versus total tissue area) was calculated for downstream statistical analysis.

**Spatial transcriptomics (Visium HD)**

Tumors from MC38 tumor-bearing mice were collected at ZT2 or ZT18 and fixed in 4% PFA and paraffin embedded (FFPE) as above. 5-µm thick sections were cut from FFPE tissue blocks using a microtome. The FFPE fixed tissue sections were placed on Schott Nexterion H glass histology slides (ThermoFisher Scientific) for deparaffinization, H&E staining and imaging according to the standard Visium HD FFPE Tissue Preparation protocol (CG000684). The mouse reagents (Visium HD, mouse transcriptome kit, 6.5mm, #1000667) were used to perform hybridization, probe amplification, transfer to Visium HD slides using the CytAssist system (10x Genomics), and for library generation according to the standard Visium HD Spatial Gene Expression Reagent Kits protocol (CG000685). Libraries were sequenced on an Illumina NovaSeq 6000 sequencer with paired end reads.

**Spatial Transcriptomics (Visium HD) Analysis**

We used Space Ranger (v3.0) to map FASTQ files to the mouse reference genome (UCSC mm10), detect the tissue section, align the sequencing data to the microscopic H&E image and the CytAssist image, and output gene-barcode matrices for further analysis. For downstream spatial analysis, we used Loupe Browser v.8 (10x Genomics) to visualize gene expression across both tissue sections. Low-expression spots were filtered to remove background noise. Spot-level clustering was performed based on global gene expression profiles, and immune and non-immune clusters were identified using canonical marker genes (See Fig. S2).

In spatial transcriptomics, calculating distances between spots or clusters using pixel coordinates is widely used (e.g SpaGCN) (*49-52*). We used a similar approach to assess spatial proximity between immune cell populations based off pixel distance. We calculated the distance between each T cell cluster to the nearest 100 myeloid cell clusters and averaged these distances to obtain a representative T cell to myeloid cell distance (See Fig. 2C). This approach limits noise from distant cells that are less likely to exert biological influence and prioritizes interactions within a plausible functional range. Distance distributions were visualized using violin plots.

To investigate the functional significance of transcriptional differences across different time points, we performed differential expression analysis between ZT2 and ZT18 at the whole-slide level. Differentially expressed genes (DEGs) were identified using the Wilcoxon rank-sum test (implemented in Scanpy v1.9.3**),** with significance thresholds set at an adjusted p-value $< 0.05$ and log2 fold-change $> 1$.

Enrichment analysis was then conducted using the gseapy (v0.10.8) Python package, employing the Enrichr framework with the Gene Ontology Biological Process 2021 (mouse) reference dataset. Statistical significance of pathway enrichment was evaluated using Fisher's exact test with Benjamini–Hochberg correction for multiple testing. Results were visualized as bar plots of –log10 adjusted p-values using Matplotlib v3.8.

To further explore the relationship between spatial proximity and immune cell signaling, we divided the range of T cell–to–myeloid cell distances into ten quantiles, from the closest (quantile 1) to the farthest (quantile 10). For each distance group, mean expression values of selected chemokine and cytokine genes were calculated across T cell and myeloid cell quantiles.

To allow comparability across genes, expression values were min–max normalized on a per-gene basis. The resulting distance-stratified expression profiles were visualized as heatmaps, with rows representing distance quantiles and columns representing genes (Fig. 2D). All analyses were performed using Loupe Browser v8 and custom Python scripts implemented in Scanpy v1.9.3, Seaborn v0.12, and Matplotlib v3.8.

For visualization of effector cytokine and chemokine expression differences, we generated dot plots comparing ZT2 and ZT18 immune clusters. Average gene expression values were computed at the cluster level, and expression fractions were calculated as the proportion of cells with nonzero expression for each gene. Dot plots were generated using Scanpy v1.9.3, with dot size encoding the fraction of expressing cells and color intensity representing the standardized average expression per cluster (min–max normalized across genes).

**FACS isolation of tumor-infiltrating DCs and bulk RNA sequencing**

To obtain tumor-infiltrating DCs, tumors were collected from MC38 tumor-bearing mice 10 days after tumor inoculation at indicated time points (ZT2 or ZT18). Single cell suspensions were obtained as previously described. Given the rarity of this cell population, single cell suspensions from five mice (n = 5 per biological replicate) were pooled per one sample sent for sequencing (total of n =15 mice or 3 biological replicates per ZT). Prior to FACS, immune cells were further purified using $CD45^+$ TIL microbeads and magnetic-activated cell sorting (Miltenyi $CD45^+$ TIL kit) according to manufacturer's instructions. $CD45^+Lineage^-(CD3^-NK1.1^-CD19^-Ly6C^-Ly6G^-F4/80^-)$CD11c+ dendritic cells were isolated via FACS using a FACSymphony S6 cell sorter (BD Biosciences). Cells were immediately immersed in Trizol LS reagent (ThermoFisher #10296010), flash frozen in liquid nitrogen and stored at -80°C to preserve RNA quality. RNA was extracted using the standard TRIzol phenol-chloroform extraction protocol and a RNeasy Micro Plus Kit (Qiagen). Briefly, a phenol chloroform extraction was performed for each sample according to the standard TRIzol LS protocol. The aqueous layer was carefully aspirated and added to an equal volume of 70% ethanol and directly added to a RNeasy spin column. RNA isolation then proceeded via the RNeasy Micro Plus Kit protocol and eluted into nuclease free water. RNA integrity and quality was accessed using a Bioanalyzer Pico Kit (Agilent Technologies) and only samples with RIN >9.0 were used for library preparation and sequencing. At least 50ng of total RNA was used per each cDNA library preparation. cDNA libraries were prepared by the UT Southwestern Genomics Core using an Illumina TruSeq mRNA library prep kit according to manufacturer's instructions. Libraries were sequenced on an Illumina Novaseq 6000 using paired end reads.

**RNA Sequencing analysis**

Reads were aligned to the mouse reference genome (UCSC mm10). Trim Galore was used for adapter and quality trimming. The mouse reference genome (mm10) and its gene annotation were downloaded from the UCSC Genome Browser and the NCBI RefSeq database. Library quality was assessed by mapping reads to mouse transcript and ribosomal-RNA sequences with the Burrows-Wheeler Aligner (BWA v0.7.17). Reads were aligned to the mouse genome with STAR (v2.7.10b), and the alignments were sorted with SAMtools (v1.16.1). Gene-level counts were obtained with the HTSeq Python package. Read counts were normalized and differentially expressed (DE) genes were identified with the DESeq2 package in Bioconductor. KEGG pathway data were retrieved via the KEGG API (https://www.kegg.jp/kegg/rest/keggapi.html). Enrichment of DE genes in KEGG pathways was evaluated with Fisher's exact test in R.

**Bulk Cytokine/Chemokine analysis from whole tumor lysates**

Tumors of equivalent weight were harvested from MC38 tumor-bearing mice at specified time points (ZT2, ZT6, ZT14, ZT18). They were weighed and added to 400 µl of lysis buffer 2 (R&D Systems) supplemented with protease and phosphatase inhibitor tablets (Roche). Tumor tissue lysates were prepared using a tissue homogenizer (Omni Tissue Master 125) and incubated on a rocker at 4°C for 30 minutes. Samples were centrifuged at 15,000 RPM for 15 minutes at 4°C. The supernatant containing cytokines/chemokines was collected and stored in multiple aliquots at -80°C. Quantification of cytokines and chemokines was performed using the Mouse Luminex Discovery Assay (R&D systems, cat# LXSAMSM) according to manufacturer's instructions on a Bio-Plex 200 (Bio-Rad). A separate aliquot was used and ran according to manufacturer's instructions for quantification of CXCL9 (Mouse ProcartaPlex Simplex Kit, Thermo-Fisher #EPX010-26061-901) and CX3CL1 (Mouse Miliplex Kit, Sigma-Millipore, #MECY2MAG-73K). Raw protein concentrations were normalized to tumor weight.

**Intratumoral rCX3CL1 administration**

Working solutions of rCX3CL1 (Recombinant mouse CX3CL1/Fractalkine R&D systems #458-MF-025/CF)) were made by dissolving rCX3CL1 in sterile PBS at a concentration of 2ng/µl. 50 µl of rCX3CL1 (100ng per mouse) was injected intratumorally into MC38 tumor-bearing mice using a 27g insulin syringe at indicated time points on days 8, 12 and 16 after tumor inoculation.

**Chromatin immunoprecipitation qPCR (ChIP-qPCR)**

For the generation of spleen-derived DCs, C57BL6/J mice were subcutaneously inoculated with 5 x $10^6$ B16-FLT3L cells (RRID:CVCL_IJ12) into the right inguinal flank. Inoculation with these cells leads to robust proliferation of $CD11c^+$ DCs in the spleen (*53*) . After 14 days, spleens were harvested and DCs purified using CD11c microbeads and magnetic-activated cell sorting (Miltenyi) as previously described (*9*). For ChIP, a total of 2 x $10^7$ DCs were used per sample. Briefly, the cells were fixed in PBS with 1% formaldehyde (Thermo Fisher) for 8 minutes and quenched with 2.5M of glycine on ice. Fixed cells were washed with PBS and resuspended in lysis buffer (50 mM Tris-HCl pH 7.5, 10 mM EDTA, 0.5% L-lauryl sarcosine, 1 mM PMSF, and EDTA-free protease inhibitor cocktail (Roche)) and sonicated on a Covaris S2 sonicator (14 × 30 seconds at 4°C) to shear chromatin and obtain fragments of 0.2-0.8 kilobases in size, which was verified on a DNA gel. Immunoprecipitation was performed using 30 µg of chromatin and incubated overnight with 1µg anti-BMAL1 or chicken IgY control (Abcam) antibodies. The next day 20µL of IgY precipitating agarose resin (GenScript) was added and incubated for 2 hours at 4°C with rotation. DNA was eluted into nuclease-free water and purified using a QiaQuick PCR purification kit (Qiagen). qPCR was performed using *Cx3cl1* primers (designed using NCBI Primer-BLAST) using Platinum SYBR green qPCR super mix (ThermoFisher Scientific) on a QuantStudio 7 Flex Real-Time PCR System (Applied Biosystems).

**Statistical analysis**

GraphPad Prism v.10 was used for statistical analysis. All data sets were tested for normality (e.g. Shapiro-Wilk). Data sets with normal distribution were analyzed with parametric tests, such as a Student t-test or one-way ANOVA with Bonferroni post-test. For non-normal distributions, non-parametric tests, such as the Mann-Whitney U test or Kruskal-Wallis test were applied. Survival was analyzed using the Mantel-Cox log-rank test.

**Language Editing Assistance**

OpenAI's ChatGPT (version 4.0 and 5.0) was used solely for language editing (e.g., grammar, clarity, and brevity) and for suggesting alternative phrasings during manuscript preparation. The tool was not used for study design, data generation, and analysis. It was not used to generate references. All suggestions from ChatGPT were reviewed, revised, and approved by the authors. No text was inserted verbatim without subsequent human editing. The authors take full responsibility for the content of the manuscript.

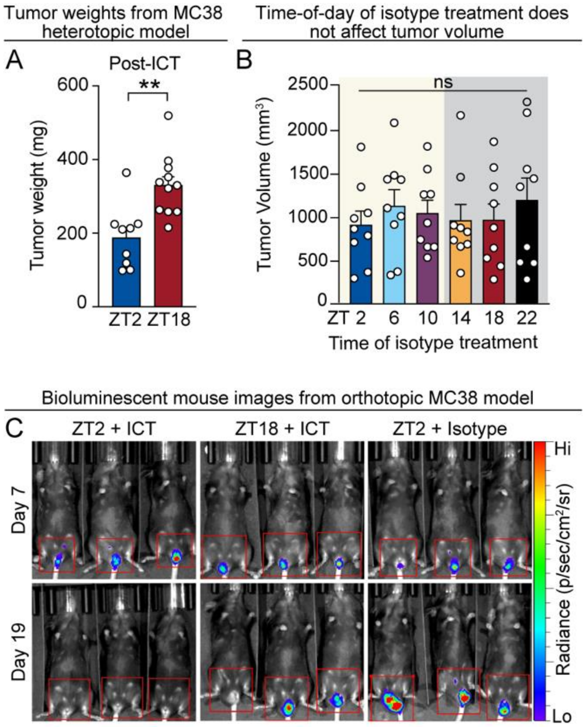


**Fig. S1. ICT efficacy is time-of-day–dependent**

**(A)** MC38 tumor weights on day 17 following anti–PD-1 treatment (ICT) at ZT2 or ZT18, n = 9–10 mice per group.

**(B)** Tumor volumes at day 20 following isotype (200 µg rat IgG2a) administered at six distinct zeitgeber times (ZT2 to ZT22); n = 9 mice per group.

**(C)** Representative bioluminescent mouse images from orthotopic MC38-Luc model on day 7 (prior to ICT or isotype treatment) and after treatment at ZT2 or ZT18. Red squares represent area (location of MC38-Luc rectal tumors) that was quantified in Fig. 1F.

ICT, immune checkpoint inhibitor therapy; ZT, zeitgeber time. Statistical analysis by Mann-Whitney test (A) and Kruskal-Wallis test (B). Dots represent results from individual mice. Bars denote mean ± SEM. All data are representative of ≥2 independent experiments. $^{**}p < 0.01$. ns, not significant.

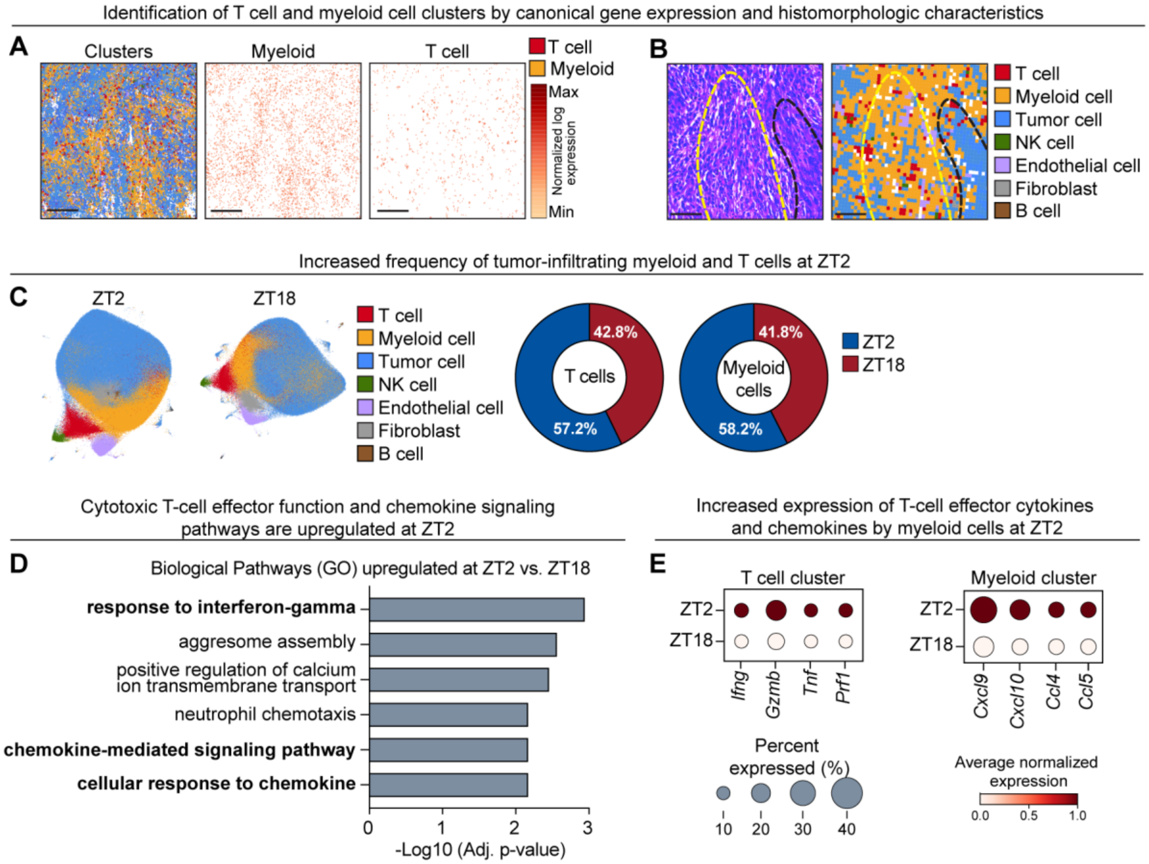


**Figure S2. Spatial transcriptomics analysis of tumor-infiltrating myeloid and T cells**

**(A)** Representative spatial transcriptomics images showing annotated cell clusters (left image), gene expression analysis for identifying myeloid cells (center image) and T cells (right image). Myeloid cell identifying genes (*Itgax, itgam, adgre1, lyz2, batf3, clec9a, csf1r*) and T cell identifying genes (*Cd3e, cd3d, cd8a, cd8b1, cd4, cd3g)* are shown. Black scale bar represents 200 μm.

**(B)** Identification of cell clusters by histomorphologic characteristics. H&E image (left) and spatial transcriptomics images (right). Yellow dotted line highlighting immune cell-rich area and black dotted line indicating tumor cell-rich area. Black scale bar represents 100 μm.

**(C)** UMAP plots of all annotated clusters identified at ZT2 and ZT18 (left). Proportional contribution of each immune cell type at ZT2 and ZT18 (right).

**(D)** Gene ontology pathway analysis of the top biological pathways upregulated at ZT2 vs. ZT18.

**(E)** Average gene expression levels of selected chemokines in myeloid cell clusters and effector cytokines in T cell clusters at ZT2 versus ZT18.

NK, natural killer cell; ZT, zeitgeber time; GO, gene ontology; UMAP, Uniform Manifold and Approximation Plot.

Key diffferentially expressed transcripts used to identify single cell-RNAseq clusters.

**A**

| Cluster name | Key differentially expressed transcripts to identify clusters |
|---|---|
| 1 B cells | *CD79a, CD79b, Mzb1, Ms4a1, Cd19* |
| 2 Tregs | *Foxp3, Icos, Ikzf2, Izumo1r, Tnfrsf4, Ctla4* |
| 3 Plasmacytoid Dendritic Cells (pDCs) | *Siglech, Gm21762, Cd300c* |
| 4 CD8+ T cells | *Cd8b1, Cd8a, Pdcd1, Tox* |
| 5 CD4+ T cells | *Cd4, Bcl11b, Tcf7, Dapl1* |
| 6 Gamma Delta T cells | *Tcrg-C1, Trdc, Tcrg-C4, Il17a* |
| 7 Macrophages/Monocytes | *Itgam, Adgre1, Ly6c2, Csf1r, C1qc, C3, Arg1, Lyz2, H2-Eb1, H2-Ab1* |
| 8 PMN-MDSCs | *S100a8, S100a9, Cxcr2, Csf3r* |
| 9 Ki-67+ Cells | *Mki67* |
| 10 NK Cells | *Klre1, Gzma, Gzmc, Ncr1, Klra8,* |
| 11 Conventional Dendritic Cells (cDCs) | *Itgax, Clec9a, Batf3, Flt3, H2-Aa, H2-Eb1, Cd209a* |

Identification of CD8+ T cell and dendritic cell clusters by canonical gene expression

**B**

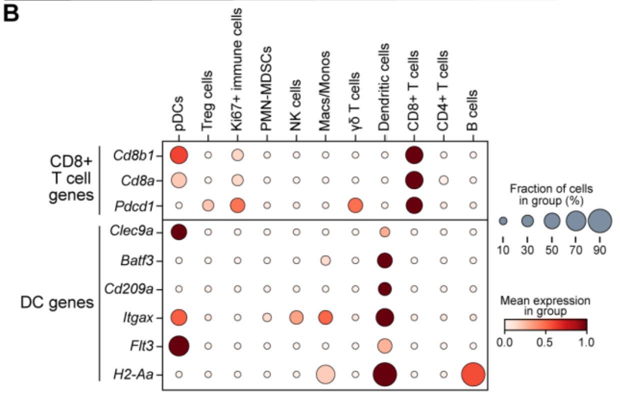


Key CD8+ T cell and dendritic cell identifying genes are highly expressed in the expected clusters.

**C**

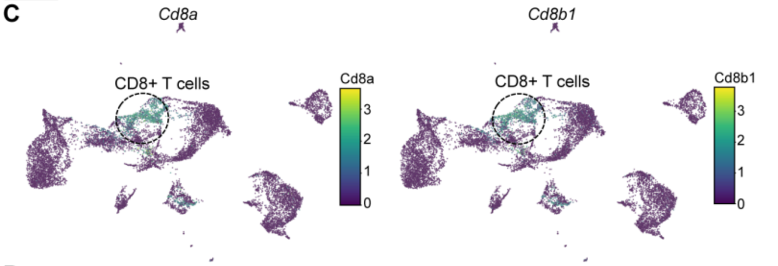


**D**

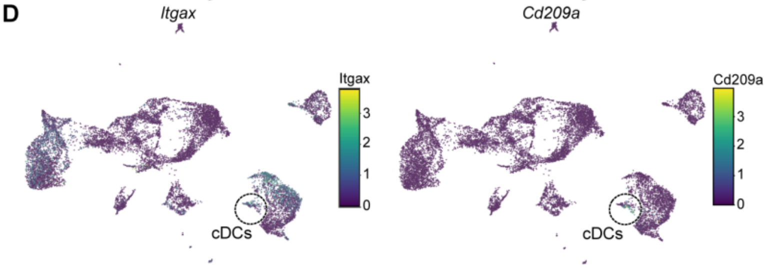


Increased proportions of PMN-MDSCs and pDCs at ZT18

**E**

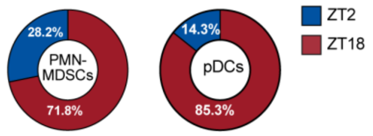


Myeloid cells have increased expression of anti-tumor chemokines at ZT2

**F**

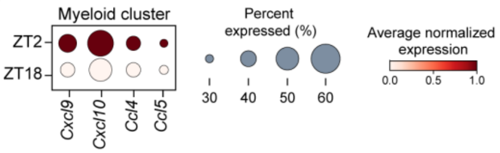

**Figure S3. scRNA-seq analysis of tumor-infiltrating leukocytes**

**(A)** Table with the list of differentially expressed transcripts used to identify immune cell clusters in UMAPs from Fig. 2G.

**(B)** Average expression of key differentially expressed transcripts used to identify $CD8^+$ T cell and cDC clusters in the scRNA-sequencing datasets.

**(C)** UMAP plots highlighting expression of representative $CD8^+$ T-cell identifying transcripts (e.g. *cd8a, cd8b1).*

**(D)** UMAP plots highlighting expression of representative cDC identifying transcripts (e.g. *itgax, cd209a*).

**(E)** Proportional contribution of PMN-MDSCs and pDCs at ZT2 versus ZT18.

**(F)** Average expression of chemokines in myeloid cell clusters (monocyte/macrophage, cDCs) between ZT2 and ZT18.

scRNA-seq, single cell RNA sequencing; ZT, zeitgeber time; Treg, T regulatory cells, PMN-MDSC, polymorphonuclear myeloid derived suppressor cell; NK cells, natural killer cells; DC, dendritic cells; pDC, plasmacytoid dendritic cell; UMAP, uniform manifold approximation and projection.

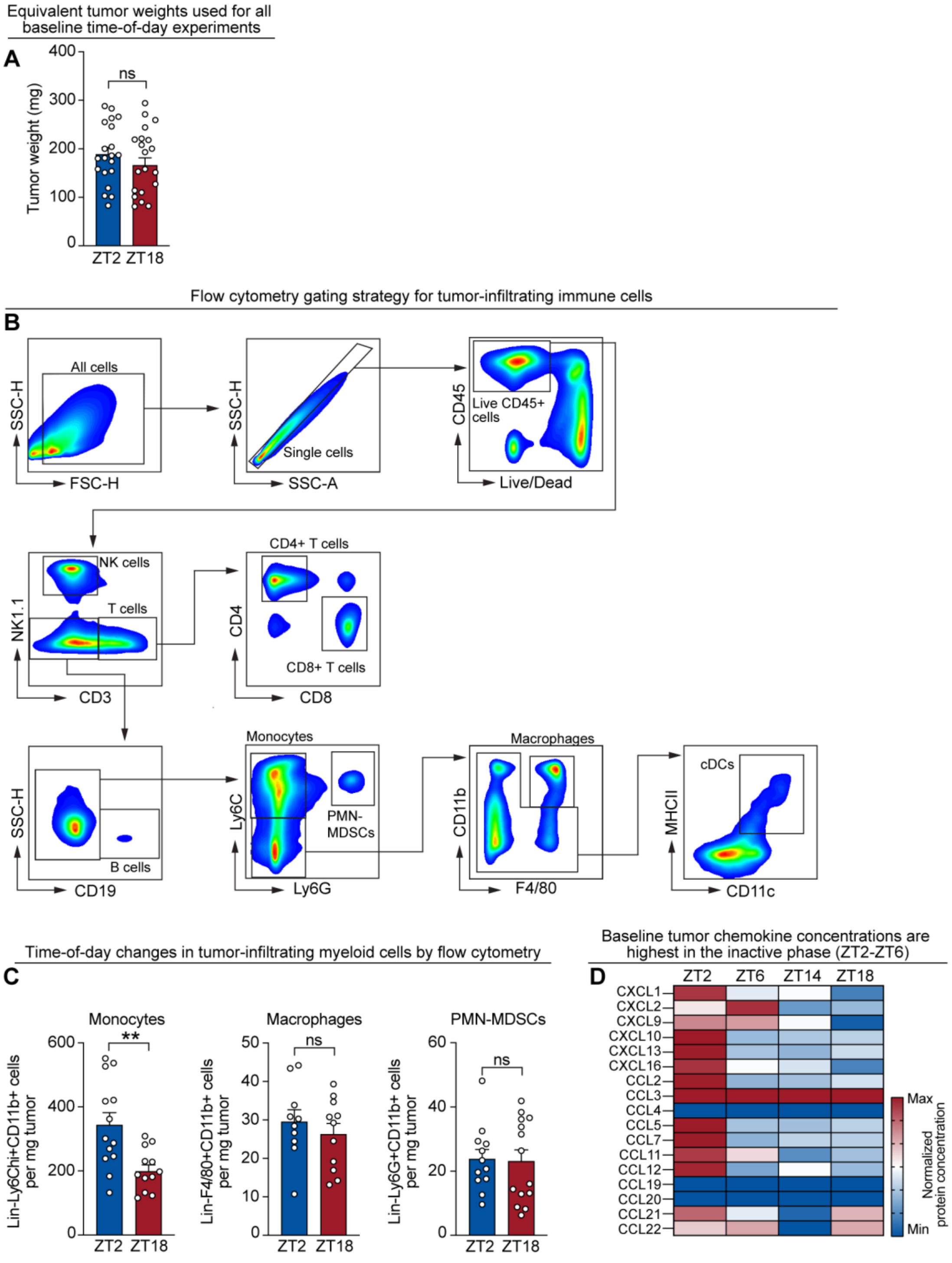


**Fig. S4. Time-of-day fluctuations in tumor-infiltrating myeloid cells and chemokines**

**(A)** Representative MC38 tumor weights on day 10 prior to subsequent immune profiling experiments (flow cytometry, IHC, Luminex) from 3 independent experiments, n = 20 mice per group.

**(B)** Flow cytometry gating strategy for tumor-infiltrating immune cells.

**(C)** Quantification of tumor-infiltrating monocytes, macrophages and PMN-MDSCs at ZT2 and ZT18 by flow cytometry; n = 10–13 mice per group.

**(D)** Heatmap representing protein concentrations of total chemokine levels from tumor lysates generated at ZT2, ZT6, ZT14, or ZT18 as measured by multiplex Luminex assay; Data represents mean protein concentration from $n = 8–10$ mice per group. Data was averaged and scaled.

ZT, zeitgeber time; PMN-MDSC, polymorphonuclear myeloid derived suppressor cell. For each experiment, points represent results from individual mice. Bars denote mean ± SEM. All data are representative of ≥2 independent experiments. Statistical analysis by Students t-test in (A) and Mann-Whitney test in (C). $^{**}p < 0.01$. ns, not significant.

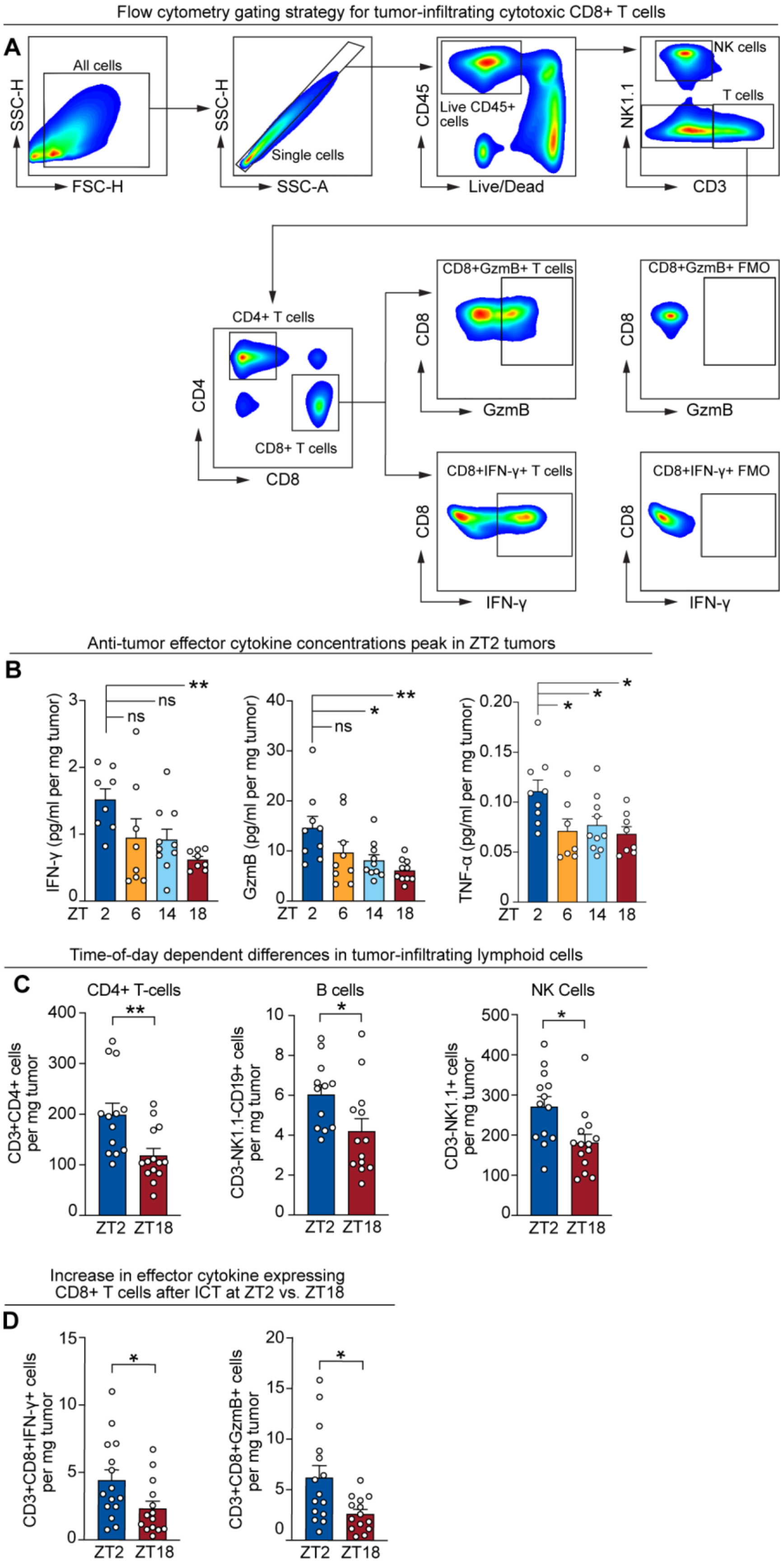
Flow cytometry gating strategy for tumor-infiltrating cytotoxic CD8+ T cells
A
All cells
SSC-H
FSC-H
Single cells
SSC-A
CD45
Live CD45+ cells
Live/Dead
NK cells
T cells
NK1.1
CD3
CD4+ T cells
CD8+ T cells
CD4
CD8
CD8+GzmB+ T cells
GzmB
CD8+GzmB+ FMO
CD8+IFN-γ+ T cells
IFN-γ
CD8+IFN-γ+ FMO
Anti-tumor effector cytokine concentrations peak in ZT2 tumors
B
IFN-γ (pg/ml per mg tumor)
GzmB (pg/ml per mg tumor)
TNF-α (pg/ml per mg tumor)
ns
*
**
ZT 2 6 14 18
Time-of-day dependent differences in tumor-infiltrating lymphoid cells
C
CD4+ T-cells
B cells
NK Cells
CD3+CD4+ cells per mg tumor
CD3-NK1.1-CD19+ cells per mg tumor
CD3-NK1.1+ cells per mg tumor
ZT2 ZT18
Increase in effector cytokine expressing CD8+ T cells after ICT at ZT2 vs. ZT18
D
CD3+CD8+IFN-γ+ cells per mg tumor
CD3+CD8+GzmB+ cells per mg tumor
ZT2 ZT18

**Figure S5. Time-of-day regulation of tumor-infiltrating cytotoxic CD8+ T cells**

**(A)** Flow cytometry gating strategy for assessing tumor-infiltrating T cells, including cytotoxic interferon-$\gamma^+$ or granzyme $B^+$ $CD8^+$ T cells with FMO controls.

**(B)** Quantification of IFN-γ, granzyme B, and TNF-α protein levels in tumor lysates collected at ZT2, ZT6, ZT14 and ZT18, measured by Luminex assay; n = 8–10 mice per group.

**(C)** Quantification of tumor-infiltrating $CD4^+$ T cells, B cells and NK cells at ZT2 and ZT18 by flow cytometry; n = 10–13 mice per group.

**(D)** Quantification of IFN-$\gamma^+$ (left) and granzyme $B^+$ (right) $CD8^+$ T cells in tumors after ICT at ZT2 versus ZT18 on day 17 after three doses of ICT; n =14–15 mice per group.

FMO, fluorescence minus one control; ZT, zeitgeber time; ICT, immune checkpoint inhibitor therapy; NK, natural killer cells. Statistical tests: One-way ANOVA (B), Mann-Whitney test (C, D). For each experiment, points represent results from individual mice. Bars denote mean ± SEM. All data are representative of ≥2 independent experiments. *$p < 0.05$; **$p < 0.01$. ns, not significant.

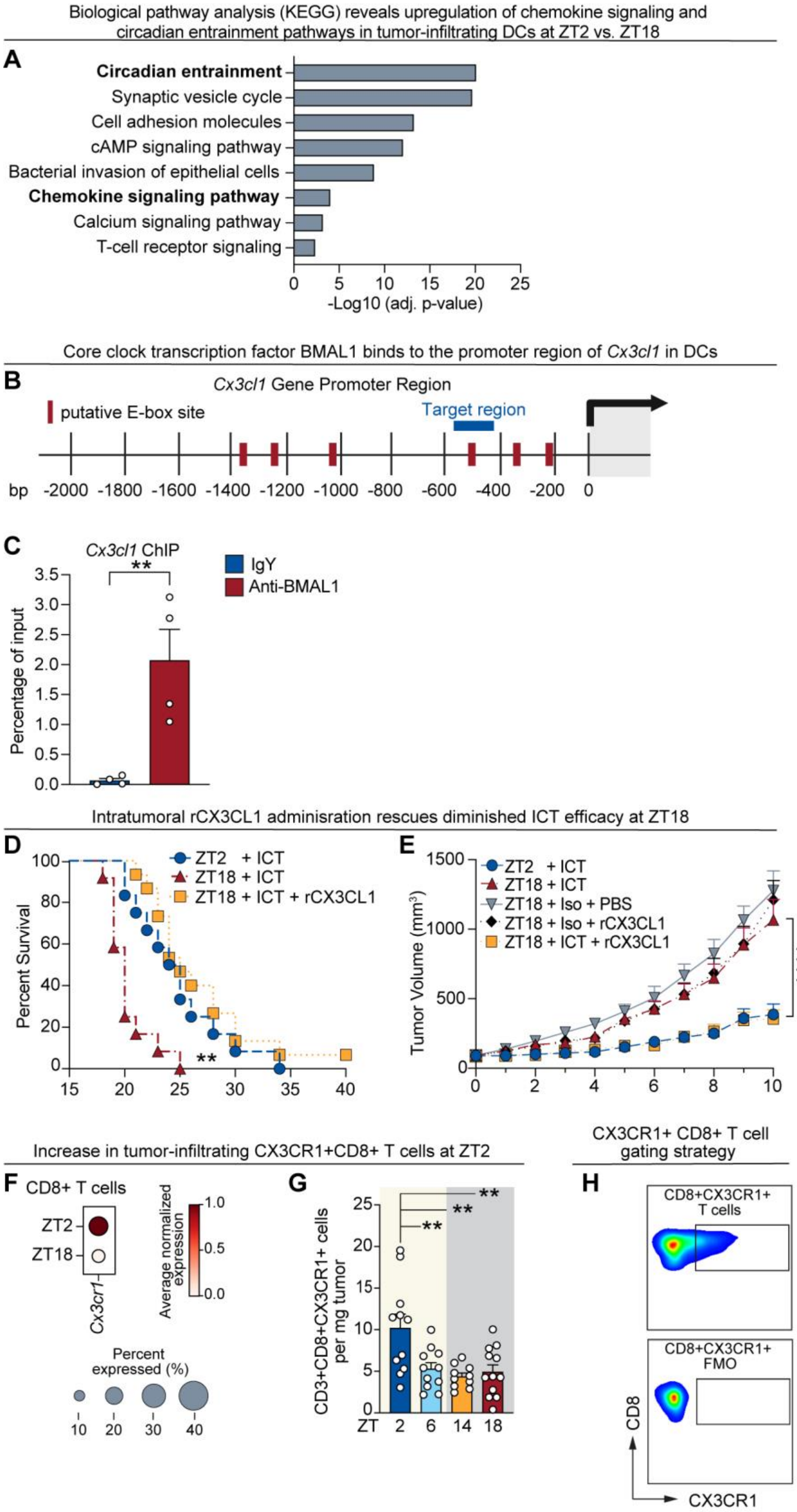
Biological pathway analysis (KEGG) reveals upregulation of chemokine signaling and circadian entrainment pathways in tumor-infiltrating DCs at ZT2 vs. ZT18
A
Circadian entrainment
Synaptic vesicle cycle
Cell adhesion molecules
cAMP signaling pathway
Bacterial invasion of epithelial cells
Chemokine signaling pathway
Calcium signaling pathway
T-cell receptor signaling
0 5 10 15 20 25
-Log10 (adj. p-value)
Core clock transcription factor BMAL1 binds to the promoter region of Cx3cl1 in DCs
B
Cx3cl1 Gene Promoter Region
putative E-box site
Target region
bp -2000 -1800 -1600 -1400 -1200 -1000 -800 -600 -400 -200 0
C
Cx3cl1 ChIP
IgY
Anti-BMAL1
Percentage of input
**
Intratumoral rCX3CL1 adminisration rescues diminished ICT efficacy at ZT18
D
ZT2 + ICT
ZT18 + ICT
ZT18 + ICT + rCX3CL1
Percent Survival
**
E
ZT2 + ICT
ZT18 + ICT
ZT18 + Iso + PBS
ZT18 + Iso + rCX3CL1
ZT18 + ICT + rCX3CL1
Tumor Volume (mm3)
****
Increase in tumor-infiltrating CX3CR1+CD8+ T cells at ZT2
F
CD8+ T cells
ZT2
ZT18
Cx3cr1
Average normalized expression
Percent expressed (%)
10 20 30 40
G
CD3+CD8+CX3CR1+ cells per mg tumor
ZT 2 6 14 18
**
CX3CR1+ CD8+ T cell gating strategy
H
CD8+CX3CR1+ T cells
CD8+CX3CR1+ FMO
CD8
CX3CR1

**Fig. S6. Circadian regulation of CX3CL1 signaling in DCs drives CX3CR1$^{+}$CD8$^{+}$ T cell recruitment**

**(A)** KEGG pathway analysis from tumor-infiltrating dendritic cell RNA-seq experiment from Fig. 5A.

**(B)** Gene map of the promoter region of *Cx3cl1*. Red lines indicate putative BMAL1 binding sites called E-box binding motifs (CANNTG, NN = any nucleotide). Blue line indicates target region validated in ChIP-qPCR experiment in S6C. 0 = start codon.

**(C)** CD11c$^{+}$ DCs were purified from the spleens of C57BL/6 mice. 2 x $10^7$ DCs per sample underwent ChIP-qPCR to confirm anti-BMAL1 vs. IgY binding to the E-box sites in the promoter region of *Cx3cl1*. n = 4 samples per group.

**(D)** Survival of C57BL6/J mice treated with ICT (anti–PD-1) ± intratumoral rCX3CL1 at ZT2 or ZT18 (related to Fig. 5F-G) n = 13–14 mice per group.

**(E)** Tumor volume of C57BL6/J mice treated with ICT (anti–PD-1) or isotype ± intratumoral rCX3CL1 or vehicle (PBS) at ZT2 or ZT18 (related to Fig. 5F-G) n = 13–14 mice per group.

**(F)** Average e pression levels of *Cx3cr1* in the CD8$^{+}$ T-cell cluster between ZT2 and ZT18 from scRNA-seq experiment (See Fig. 2F-G).

**(G)** Quantification of tumor-infiltrating CX3CR1$^{+}$CD8$^{+}$ T cells at ZT2, ZT6, ZT14, ZT18 in C57BL/6 mice prior to ICT; n = 10–11 mice per group.

**(H)** Flow cytometry gating strategy for tumor-infiltrating CX3CR1$^{+}$ CD8$^{+}$ T cells with FMO controls.

KEGG, Kyoto Encyclopedia of Genes and Genomes; bp, base pairs; E-box, enhancer box region; ZT, zeitgeber time; ICT, immune checkpoint inhibitor therapy; rCX3CL1, recombinant CX3CL1; Iso, isotype; FMO, fluorescence minus one gating control. Statistical analysis by: Students t-test (C), Log rank test (D), Mann-Whitney test (E), Kruskal-Wallis test (G). For each experiment, points represent results from individual mice. Bars denote mean ± SEM. All data are representative of ≥2 independent experiments. *p < 0.05; **p < 0.01; ***p < 0.001; ****p < 0.0001.

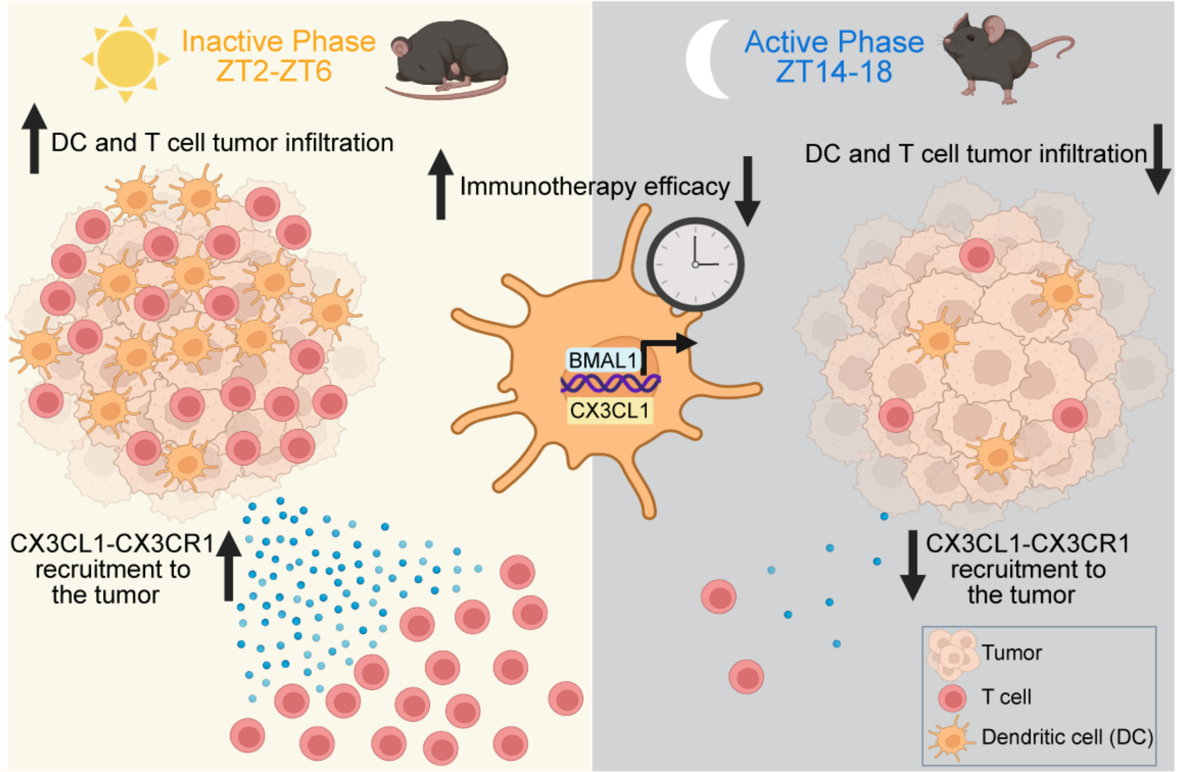


**Fig. S7. Proposed model of clock-dependent regulation of immunotherapy efficacy.**
The dendritic cell-intrinsic clock drives transcription of *Cx3cl1*, which recruits CX3CR1$^{+}$CD8$^{+}$ T cells to the tumor immune microenvironment during the inactive/light phase (ZT2–ZT6). ICT administered at this time of peak immune infiltration and activation enhances anti-tumor immunity and promotes a sustained therapeutic response. DC, dendritic cell; ZT, zeitgeber time.

**Table S1. Key Resources Table**

| Reagent or Resource | Source | Identifier |
|---|---|---|
| **Flow cytometry Antibodies** | | |
| Anti-mouse CD45 (BV650) | Biolegend | Cat# 103151 |
| Anti-mouse CD3 (BV785) | Biolegend | Cat# 100222 |
| Anti-mouse CD4 (PE-Cy5) | Biolegend | Cat# 100551 |
| Anti-mouse CD19 (BV605) | Biolegend | Cat# 115540 |
| Anti-mouse CD11c (FITC) | Biolegend | Cat# 117339 |
| Anti-mouse I-A/I-E (AF700) | Biolegend | Cat# 107622 |
| Anti-mouse CD8a (BV510) | Biolegend | Cat# 100712 |
| Anti-mouse NK1.1 (Pacific blue) | Biolegend | Cat# 100752 |
| Anti-mouse Ly6C (BV711) | Biolegend | Cat# 128037 |
| Anti-mouse Ly6G (PE-Cy7) | Biolegend | Cat# 100722 |
| Anti-mouse F4/80 (APC) | Biolegend | Cat# 123116 |
| Anti-mouse CD11b (PE-Dazzle 594) | Biolegend | Cat# 101256 |
| Anti-mouse interferon-y (APC) | Biolegend | Cat# 505810 |
| Anti-mouse granzyme B (BV421) | Biolegend | Cat# 396414 |
| Anti-mouse CX3CR1 (APC) | Biolegend | Cat# 149008 |
| Anti-mouse CD16/CD32 (Fcγ receptor) | Biolegend | Cat# 10132 |
| Ghost Dye Red 780 | Thermo Fisher Scientific | Cat# 50-105-2988 |
| **Cancer Cell Lines** | | |
| B16-F10 | ATCC | RRID:CVC_0159 |
| MC38 | Dr. Todd Aquilera | RRID:CVCL_B288 |
| MC38-Luciferase | ALSTEM Bio | RRID:CVCL_C8VZ |
| B16-FLT3L | Dr. Chandrasakaren Pasare | RRID:CVCL_IJ12 |
| **Chemicals, peptides and recombinant proteins** | | |
| RNAseZap | ThermoFisher Scientific | Cat# AM9780 |
| 4% Paraformaldehyde (PFA) in PBS | ThermoFisher Scientific | Cat# J19943-K2 |
| RBC lysis buffer | Invitrogen | Cat# 00433357 |
| Fetal Bovine Serum (Heat inactivated) | Gibco | Cat# 10082147 |
| DMEM | Gibco | Cat# 11995-065 |
| Recombinant CX3CL1 | R&D Systems | Cat# 458-MF-025/CF |
| Recombinant murine GM-CSF | R&D Systems | Cat# 415-ML-020/CF |
| Recombinant murine IL-4 | R&D Systems | Cat# 404-ML-025/CF |
| Cultrex Basement Membrane Extract, Type 3, Pathclear | R&D Systems | Cat# 3632-010-02 |
| D-Luciferin, potassium salt (endotoxin free) | Gold Biotechnology | Cat# eLUCK-1G |
| RPMI 1640 | ATCC | Cat# 20-3001 |
| Cell Activation Cocktail (with Brefeldin A) | Biolegend | Cat# 423304 |
| BD Horizon Brilliant Stain Buffer Plus | BD Biosciences | Cat# 566385 |
| Platinum SYBR green qPCR super mix | ThermoFischer Scientific | Cat# 11733046 |
| **Antibodies (other than for flow cytometry)** | | |
| Anti-PD-1 (Clone RMP-1, CD279) | BioXcell | Cat# BP0146 |

| | | |
|---|---|---|
| Anti-CTLA-4 (9D9, CD152) | BioXcell | Cat# BP0164 |
| Anti-rat IgG2a, κ (isotype for anti-PD-1) | BioXcell | Cat# BE0089 |
| Anti-mouse IgG2b, κ (isotype for anti-CTLA-4) | BioXcell | Cat# BE0086 |
| Anti-CD8a (rabbit clone EPR21769) for IHC | Abcam | Cat# ab217344 |
| Anti-CD11c (rabbit D1V9Y) for IHC | Cell Signaling | Cat# 39143SF |
| Anti-BMAL1 (for ChIP) | Dr. Joseph Takahashi | Koike et. al 2012 (*54*)<br>Yoo et. al 2013 (*55*) |
| Anti-IgY (for ChIP) | Abcam | Cat# ab50579 |
| **Critical commercial assays** | | |
| Tumor dissociation kit, mouse | Milentyi Biotec | Cat # 23225 |
| CD45 (TIL) Microbeads, mouse | Miltenyi Biotec | Cat# 130-110-618 |
| CD11c Microbeads Ultrapure, mouse | Miltenyi Biotec | Cat# 130-125-835 |
| LS Columns | Miltentyi Biotec | Cat# 130-042-401 |
| Cyto-fast Fix-Perm Buffer Set | Biolegend | Cat# 426803 |
| Chromium Single Cell 3' v3.1 Gene Expression Kit | 10x Genomics | Cat# PN-1000268 |
| ImPRESS Excel Amplified Polymer Staining Kit, Anti-Rabbit IgG, Peroxidase. | Vector Laboratories | Cat# MP-7601-50 |
| Visium HD, mouse transcriptome kit, 6.5mm, #1000667 | 10x Genomics | Cat#1000667 |
| RNeasy Plus Micro Kit | QIAGEN | Cat# 74034 |
| TRIzol LS | ThermoFisher Scientific | Cat# 10296028 |
| Lysis buffer 2 | R&D systems | Cat# 89534 |
| Mouse Luminex Discovery Assay | R&D Systems | Cat# LXSAMSM |
| Mouse Miliplex Kit, CX3CL1 | Sigma-Millipore | Cat# MECY2MAG-73K |
| Mouse ProcartaPlex Simplex Kit, CXCL9 | ThermoFisher Scientific | Cat# EPX010-26061-901 |
| QiaQuick PCR purification kit | Qiagen | Cat# 28104 |
| **Experimental mouse models** | | |
| C57BL/6J | Jackson Laboratory | Stock No. 000664 |
| *Bmal1*$^{flox/flox}$ | Jackson Laboratory | Stock No. 007668 |
| CD11c-Cre (i.e *Itgax*-cre) | Jackson Laboratory | Stock No. 008068 |
| CD8a-Cre | Jackson Laboratory | Stock No. 008766 |
| *Cx3cr1*$^{-/-}$ | Jackson Laboratory | Stock No. 005582 |
| **Deposited Data** | | |
| Spatial transcriptomics data | This paper | |
| Single cell RNA-seq data | This paper | |
| RNA Seq data | This paper | |
| **Software and algorithms** | | |
| Fiji (ImageJ) | Fiji (ImageJ) | RRID:SCR_002285 |
| GraphPad PRISM v.10.0 | GraphPad Software | RRID:SCR_002798 |

| | | |
|---|---|---|
| NovoExpress v.1.6.1 | Agilent | RRID:SCR_024676 |
| Loupe Browser v.8.0 | 10x Genomics | RRID:SCR_018555 |
| Trim Galore | https://www.bioinformatics.babraham.ac.ukprojects/trim_galore/ | RRID:SCR_011847 |
| STAR v2.7.10b | https://code.google.com/archive/p/rna-star/ | RRID:SCR_004463 |
| BWA-MEM v0.7.17 | https://github.com/bwa-mem2/bwa-mem2 | RRID:SCR_022192 |
| SAMtools v1.16.1 | https://github.com/samtools/samtools/releases/ | RRID:SCR_002105 |
| HTSeq v2.0.5 | https://htseq.readthedocs.io/en/latest/ | RRID:SCR_005514 |
| DESeq2 | https://bioconductor.org/packages/devel/bioc/vignettes/DESeq2/inst/doc/DESeq2.html | RRID:SCR_015687 |
| LivingImage Software | PerkinElmer | RRID:SCR_014247 |
| Cell Ranger Software V6.0 | 10x Genomics | RRID:SCR_017344 |
| Seurat v5. in R | https://satijalab.org/seurat/articles/get_started.html | RRID:SCR_016341 |
| ImageScope v12.3 | Leica Biosystems | RRID:SCR_020993 |
| Space Ranger v3.0 | 10x Genomics | RRID:SCR_025858 |
| gseapy v0.10.8 | https://gseapy.readthedocs.io/en/latest/ | RRID:SCR_025803 |
| Scanpy v1.9.3 | https://scanpy.readthedocs.io/en/stable/ | RRID:SCR_018139 |
| Matplotlib v3.8 | https://matplotlib.org | RRID:SCR_008624 |
| Seaborn v0.12 | https://seaborn.pydata.org | RRID:SCR_018132 |
| UCSC Genome Browser | https://genome.ucsc.edu | RRID:SCR_005780 |
| **Key Equipment** | | |
| QuantStudio 7 Flex Real-Time PCR System | Applied Biosystems | Cat# 4485701 |
| IVIS Spectrum Imager | PerkinElmer | Cat# 124262 |
| Novocyte Advanteon Flow Cytometer | Agilent | Cat# 2010201 |
| GentleMACS Octo Dissociator with Heaters | Miltenyi Biotec | Cat# 130-096-427 |
| Illumina NovaSeq 6000 | Illumina | Cat# 20012850 |
| Visium CytAssist | 10x Genomics | Cat# 1000441 |
| **Primers** | | |
| Primer Name | Sequence | |
| *Cx3cl1*-530 site Forward Primer | 5'-ATCGAG TGA AGC TCT GTG TG-3' | N/A |
| *Cx3cl1*-530 site Forward Primer | 5'TGG ACT CAA CAC CGA ACC T-3' | N/A |
